# Sentinels for future coral reef conditions: assessment of environmental variability and water quality in semi-enclosed inland bays in the southern Caribbean

**DOI:** 10.1101/2023.10.04.560694

**Authors:** Chiara de Jong, Iris van Os, Guadalupe Sepúlveda-Rodríguez, Milo L. de Baat, Verena Schoepf

**Author notes:** Corresponding author: [ ]. Contributed equally (Chiara de Jong, Iris van Os).

## Abstract

The mangrove-seagrass-coral reef continuum is of immense ecological and socio-economic importance, supporting biodiversity, carbon storage, coastal protection, fisheries, and tourism. The presence of extreme environmental conditions along this continuum could support adaptive refugia for climate-sensitive taxa such as reef-building corals but physicochemical conditions are rarely assessed at sufficient spatiotemporal resolution. Furthermore, coastal development and low water quality increasingly threaten these interconnected coastal ecosystems. Yet, time-integrated pollution monitoring is absent at most locations. Here, we used a multi-disciplinary approach to assess benthic cover, coral diversity, and >20 abiotic parameters characterizing two mangrove- and seagrass-dominated inland bays and two nearby coral reefs in Curaçao (southern Caribbean) during the cool, dry season and warm, wet season. This was combined with time-integrated pollution monitoring using bioindicators to assess nutrients and trace metal pollution (inland bays only), and passive samplers and bioassays to assess organic chemical pollution (all four sites) during the wet season. This approach revealed a previously undocumented extent of strong diel and seasonal environmental variability in the two inland bays, with temperature, pH and dissolved oxygen frequently reaching values predicted under moderate-to-severe future climate scenarios. In addition, the inland bays had greater nutrient concentrations (especially ammonium) and ecotoxicological risks than the nearby reefs during the wet season due to run-off, industrial- and wastewater effluents, ports and boating. Overall, our findings show that Curaçao’s inland bays have significant potential to serve as natural laboratories to study the effects of future ocean conditions on resident taxa *in situ*. This however applies within the context of strong diel fluctuations and with the caveat of co-occurring stressors. Our work confirms the important role of mangrove and seagrass habitats as resilience hotspots for climate-sensitive taxa but also highlights the urgent need to improve monitoring, water quality and protection of these valuable habitats along the mangrove-seagrass-coral reef continuum.

## Introduction

Tropical mangrove forests, seagrass beds and coral reefs are among the most diverse and productive ecosystems on the planet and can form complex cross-ecosystem connections through biological, chemical, and physical interactions (Nagelkerken et al., 2008; Ogden, 1980). Mangrove forests, for example, trap sediments and regulate carbon and nutrient cycles, while seagrass meadows play crucial roles as nurseries, thus stimulating the food web and promoting biodiversity (Alongi, 1990; McDevitt-Irwin et al., 2016). Mangroves and seagrass beds further provide biogeochemical services for corals threatened by climate change by mitigating heat-stress-induced bleaching via shading (Rogers, 2017; Yates et al., 2014). At the same time, they also generate extreme temperature, pH and oxygen environments. This may drive acclimatization or adaptation of resident corals to future ocean condition (Camp et al., 2016; Unsworth et al., 2012). Coral reefs, on the other hand, create habitat for numerous marine species (Fisher et al., 2015) and provide shoreline protection, which in turn ensures low energy currents into seagrass beds and mangrove forests (Guannel et al., 2016). Consequently, the mangrove-seagrass-coral reef continuum offers not only essential ecological services but also important socio-economic benefits, such as tourism and fisheries (Costanza et al., 1997; Spalding et al., 2017; Torre-Castro & Rönnbäck, 2004). However, despite their importance, these habitats have experienced extensive deterioration over the past century due to the combined effects of global and local stressors, including climate change, overfishing, habitat destruction, coastal development, and low water quality (e.g. Collier & Waycott, 2014; Gilman et al., 2008; Hughes et al., 2017).

Climate change impacts significantly threaten mangrove forests, seagrass beds and coral reefs. For example, sea-level rise, increased cyclonic activity and changes in precipitation are among the key threats to mangrove forests (Friess et al., 2022), whereas warming and marine heatwaves pose the greatest threat to seagrass beds (Marbà et al., 2022). Yet, coral reefs are arguably one of the ecosystems that are most vulnerable to multiple climate change stressors because marine heatwaves can lead to mass coral bleaching and mortality on regional to global scales (Eakin et al., 2019; Hughes et al., 2017), while acidification lowers the calcification rates of many coral species (Kornder et al., 2018). In addition, deoxygenation and acute low oxygen events (i.e., hypoxia, defined as <2 mg L^-1^) have recently been recognized as emerging threats to coral reefs that will be exacerbated under future ocean warming and intensifying local stressors (Altieri et al., 2017; Pezner et al., 2023). However, in recent years, coral communities have been discovered that persist under extreme environmental conditions that frequently reach or even exceed temperature, pH and oxygen levels predicted to occur under future climate scenarios (Burt et al., 2020; Camp et al., 2018). Several of these naturally extreme environments occur along the mangrove-seagrass-coral reef continuum, such as mangrove lagoons (Camp et al., 2019; Stewart et al., 2022), seagrass beds (Camp et al., 2016), and semi-enclosed lagoons/bays (Maggioni et al., 2021; Vermeij et al., 2007), and can act as natural laboratories to understand mechanisms of climate change resistance and adaptation. Yet, many sites that could act as potential natural laboratories and resilience hotspots remain undocumented due to a lack of high-resolution abiotic data. Furthermore, while these extreme environments can provide critical insights into how resident coral communities respond to future climate conditions *in situ*, the impacts of global stressors in combination with local stressors have received less attention. It is, therefore, important to identify and study natural laboratories where global and local stressors occur *in tandem* to study mechanisms of climate change resistance under more realistic multi-stressor scenarios.

With human population growth near coral reefs exceeding the global average (Wong et al., 2022), local anthropogenic stressors increasingly impact habitats along the mangrove-seagrass-coral reef continuum but are often poorly monitored. Changes in land use and coastal development result in contaminated terrestrial runoff into marine waters (Burt & Bartholomew, 2019; Fabricius, 2005; Valiela et al., 2001), which exposes tropical marine organisms to excess nutrients, sediments and chemical contaminants (Jones & Kerswell, 2003; Nalley et al., 2023; Peters et al., 1997). In addition, declines in water quality can be caused by untreated wastewater discharges, industrial activities and boating, introducing various pollutants such as pharmaceuticals, personal care products, trace metals, anti-fouling agents and oil products (Schaffelke et al., 2005). Yet, the effects of these compounds on corals, mangroves and seagrasses remain poorly understood (Lewis et al., 2011; Nalley et al., 2021, 2023). Furthermore, water quality monitoring programs are often absent or limited to spot sampling which is inadequate to capture the complex spatiotemporal dynamics of many pollutants (e.g. Den Haan et al., 2016).

These challenges can be overcome via the use of time-integrated sampling approaches. For example, within biomonitoring, seagrass or macroalgae can be used to assess nutrient and trace metal content (Fabricius et al., 2012; Govers et al., 2014a, 2014b). Similarly, passive sampling devices have been developed to assess (low) concentrations of organic chemical pollution in aquatic environments which sequester chemicals by diffusion of molecules over extended periods (Vrana et al., 2005). Comprehensive water quality assessments therefore ideally combine time-integrated pollution sampling via bioindicators or passive samplers with high-frequency measurements of physicochemical parameters such as pH and dissolved oxygen from continuous logger measurements. Extracts from passive samplers can be used for toxicity assessment, where responses of organisms (*in vivo*) or cell lines (*in vitro*) are measured in bioassays, known as effect-based methods or bioanalytical tools, to study the effects of exposure to mixtures of chemicals in the environment (De Baat et al., 2020; Neale et al., 2021).

The goal of this study was two-fold. First, we aimed to conduct a comprehensive, high-frequency assessment of key physicochemical parameters across the mangrove-seagrass-coral reef continuum to determine the full range of environmental “extremeness” during both dry and wet seasons. Temperature, light, salinity, dissolved oxygen, pH, carbonate chemistry and nutrients were measured on two coral reefs and in two adjacent, semi-enclosed inland bays that feature extensive mangrove stands and seagrass meadows. The second goal was to conduct time-integrated pollution monitoring at all four sites using both bioindicators and passive sampling to gain insight into water quality and potential ecotoxicological risks to marine life. Passive sampling was paired with a battery of two *in vivo* and two *in vitro* bioassays to assess the mixture toxicity of organic chemicals. Subsequently, effect-based trigger values were used to categorize the water quality profile across the mangrove-seagrass-coral reef continuum. This state-of-the-art monitoring approach allowed us (1) to establish an environmental baseline against which future monitoring can be benchmarked, and (2) to assess whether these inland bays could serve as previously unrecognized natural laboratories and resilience hotspots for corals under climate change. Importantly, our study provides a powerful water quality framework that can be used by natural resource managers to monitor habitats along the mangrove-seagrass-coral reef continuum and conserve their high ecological and socio-economic value (Barbier et al., 2011).

## Material and Methods

### Study sites

Four sites on the island of Curaçao, southern Caribbean, were studied in March and October/November 2020, which represent the cooler dry season and the warmer wet season, respectively. The environmental variability and water quality were measured in two inland bays, Santa Martha Bay (SMB) (12°16’27.34”N, 69°07’39.66”W) with an average depth of 3 m and Spaanse Water Bay (SWB) with an average depth of 5-12 m (12°04’15.35”N, 68°51’30.61”W), and two nearby fringing reef sites, Santa Martha Reef (SMR) (12°15’59.56”N, 69°07’39.66”W) and Director’s Bay, referred to hereafter as ‘Spaanse Water Reef’ (SWR) (12°03’53.48”N, 68°51’36.69”W) (Figure 1). The two inland bays were selected based on their comparability in geomorphology, size, proximity to and availability of a reference reef site, ease of access, and presence of (similar) coral communities, seagrass and mangroves (Debrot et al., 1998; van Duyl, 1985; Vermeij et al., 2007). The bays differ in the degree of local anthropogenic stressors. Spaanse Water Bay has experienced significant coastal development over the last decades, resulting in residential use with insufficient sewage infrastructure, causing pollution and eutrophication. In contrast, Santa Martha Bay is located in a sparsely populated area with an abandoned resort undergoing reconstruction, and is thus much less impacted by local stressors than Spaanse Water Bay.

**Figure 1.**
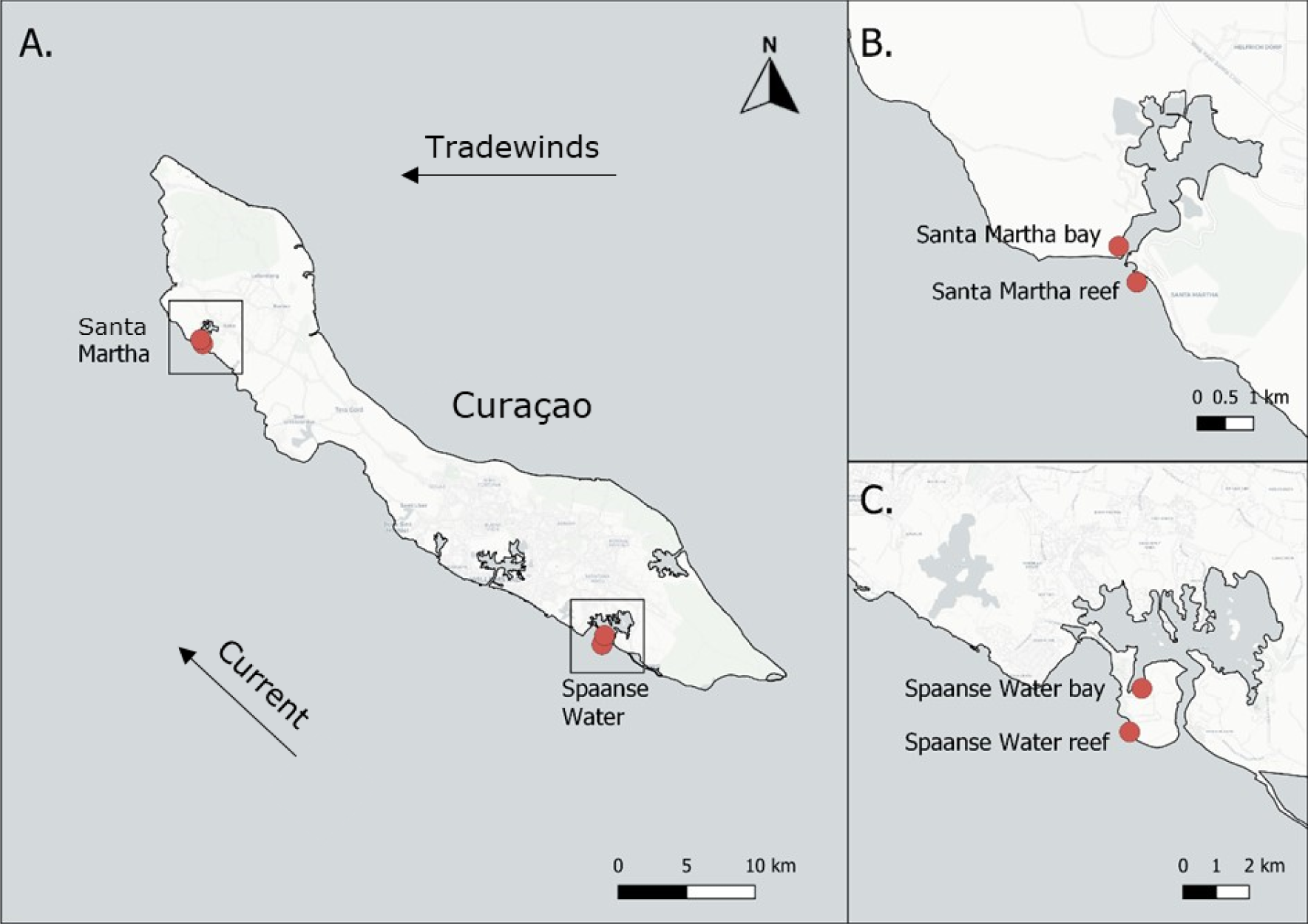
A. Map of Curaçao with sampling sites shown in red. Closer view of Santa Martha Bay and nearby fringing reef (B) and Spaanse Water Bay and nearby fringing reef (C).

### Benthic community assessment

Benthic community surveys for all sites were conducted in March 2020 using continuous line transects to characterize the immediate area surrounding the abiotic monitoring stations (see below). Given the large area of the inland bays and their heterogenous nature, the measured benthic cover is therefore only representative of the monitoring area, not the bays as a whole. In each site, six 9.9 m transects were randomly placed up to 50 m away from the cement block with environmental data loggers. Photos of the benthic community adjacent to the transect line were taken every 20 cm using a high-resolution camera (Panasonic Lumix DMC-TZ70) and the benthic cover underneath every 1 cm was analysed. A total of eleven categories were identified to describe the benthic community: hard coral, soft coral, seagrass, macroalgae, soft sediment, rubble, hard substrate, upside-down jellyfish (*Cassiopea* sp.), sea urchin (*Diadema* sp.), invertebrates and other. *Diadema* sp. and *Cassiopea* sp. were analysed as separate categories and not included in the “invertebrates” category as these organisms may have important ecological functions associated with coral development or influence physicochemical conditions (Edmunds & Carpenter, 2001; Welsh et al., 2009). Scleractinian corals were identified to species level when possible. The benthic cover was calculated according to the following formula: % benthic cover 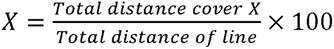

### Environmental monitoring of temperature, salinity, pH_T_, DO and PAR

Temperature, conductivity and dissolved oxygen (DO) were continuously logged, whereas pH and photosynthetically active radiation (PAR) were semi-continuous logged over two weeks during the dry season (12-03-2020 to 21-03-2020) and five weeks during the wet season (28-10-2020 to 02-12-2020) at all four sites. Logging frequency was 15 min for the conductivity and PAR loggers, and five min for the pH and DO loggers.. The combined temperature and conductivity loggers (Odyssey® Conductivity and Temperature) were calibrated with the factory-provided calibration equations using the Odyssey data logging software. pH loggers (HOBO® pH and Temperature Data Logger (MX2501)) were retrieved and calibrated every other day using TRISbuffer at two different temperatures to determine seawater pH on the total scale (pH_T_) as described in Dickson et al., (2007). TRIS buffers were purchased from A.G. Dickson. Calibration of the DO loggers (HOBO® Dissolved Oxygen Logger (U26-001)) was done by placing them in 100% saturated saltwater, whereafter they were placed in 0% saturated saltwater (obtained by adding sodium sulphite). PAR loggers (Odyssey® Xtreem Photosynthetic Active Radiation Logger) were calibrated underwater via simultaneous deployment with a Hydrolab DS5 Sonde (OTT Messtechnik GmbH & Co., Germany) equipped with a Li-Cor 4 pi spherical sensor for 12 hours (7 am – 7 pm). Loggers were placed on a cement block, ensuring that the PAR sensor and conductivity cell were vertically mounted (Figure S1a). The loggers were cleaned of debris every other day with a toothbrush. Further details are given in the Supplemental Information.

### Seawater carbonate chemistry

Discrete water samples were collected in both dry and wet seasons on multiple random days to determine seawater carbonate chemistry (via total alkalinity and pH_T_) and nutrient concentrations. Total alkalinity was analysed using potentiometric titration on a Metrohm 716 DMS Titrino (Herisau, Switzerland). Seawater pCO_2_ and aragonite saturation state (Ω_arag_) were calculated from in situ pH_T_, temperature, salinity and total alkalinity using the Carb function (flag = 8) in the seacarb package (version = 3.2.14) in R (2021.09.0).

### Nutrients and trace metals

Nitrate, nitrite, ammonium and phosphate concentrations (µmol L^-1^) were measured using a Skalar SAN^++^ system autoanalyzer. In addition, seagrass leaves were used as bioindicators for nutrient pollution (Govers et al., 2014a) and trace metal pollution (Govers et al., 2014b) because they integrate environmental conditions over time and therefore record ecologically relevant levels (Katharina E. Fabricius et al., 2012). Seagrass leaves were only collected from the two inland bay sites and only in the wet season (November 2020). Ten shoots (1 m distance between shoots) of *Thalassia testudinum* and *Halophila stipulacea* with roots were collected in Spaanse Water Bay, whereas only 10 shoots of *Halophila stipulacea* were collected in Santa Martha Bay due to the absence of *Thalassia testudinum*. The seagrass shoots were split into roots, rhizomes and leaves, after which only the leaves were used. The dried seagrass leaves were then analysed for trace metal concentrations (As, Cd, Co, Cr, Cu, Fe, Ni, P, Pb, Se, Zn) and ratios and percentages of carbon (C), nitrogen (N) and phosphate (P). Further details are given in the Supplemental Information.

### Sediment traps

Replicate sediment traps were deployed in the wet season only (November 2020) to collect sediment particles in the water column during 16 and 19 days at all four sites (see Table S1 for dates). Sediment traps were constructed following the recommendations of Storlazzi et al., (2011) for coral reef environments. To determine if trapped sediments were from local resuspension or transportation, benthic sediment at the base of each trap was also sampled. After retrieval of the sediment traps, the content was dried at 60 °C for 48 hours and analysed for weight and particle size characteristics: >1 mm, 500 to <1000 μm, 250 to <500 μm, 125 to <250 μm, 63 to <125 μm and <63 μm. The sediment trap collection rate (mg cm^2^ day^−1^) was calculated as the weight of sediment trapped (mg) divided by the number of days the trap was deployed and the surface area of the trap (cm^2^). The amount of organic matter was determined by burning the sediment samples in the muffle furnace at 450°C for 6 hours. The organic matter trap collection rate was calculated in the same way as the sediment trap collection rate. Further details are provided in the Supplemental Information.

### Effect-based chemical water quality assessment

#### Passive sampling of organic pollutants

Polar organic chemical integrative samplers (POCIS) (Alvarez et al., 2004) containing 200 mg of hydrophilic-lipophilic balance (HLB) sorbent were deployed for approximately four weeks at all four sites (see Table S1 for dates) during the wet season for the passive sampling of organic chemicals. Forty POCIS were constructed according to De Baat et al., (2020) and stored at 4°C in food-grade Mylar zip lock bags until deployment (except during transport to Curaçao). At each site, seven POCIS were attached to a stainless steel frame and retained in a stainless steel cage. The cage was placed on a cement block and deployed at the monitoring stations next to the cement block with loggers (Figure S1b). After retrieval, each POCIS was disassembled and the HLB sorbent of all 7 POCIS per site was pooled in a 50 mL Greiner tube and stored at -20 °C until extraction. The HLB sorbent was freeze-dried for 3 days at -52 °C to remove any remaining water and extracted according to De Baat et al., (2020) with slight modifications. Further details on the extraction of HLB sorbent and calculations of sampled water volumes by POCIS are provided in the Supplemental Information.

#### Bioassays

A battery of four bioassays was exposed to the POCIS extracts to determine levels of chemical pollution at the four sites (Table 1). Before application in the bioassays, the POCIS extracts were evaporated under a gentle stream of N_2_ to dryness and reconstituted in dimethyl sulfoxide (DMSO). A whole organism (*in vivo*) algal bioassay was performed with the marine alga *Dunaliella tertiolecta* at the University of Amsterdam to assess photosynthetic inhibition (Sjollema et al., 2014) using pulse amplitude modulation (PAM) fluorometry according to de Baat et al., (2018). An *in vivo* bacterial bioluminescence inhibition assay using *Alliivibrio fischeri* was performed at the Vrije Universiteit Amsterdam according to Hamers et al., (2001) to assess bacterial luminescence inhibition (commercially known as the Microtox® assay). Dose-response curves for the *in vivo* assays were made in GraphPad Prism (GraphPad Software Inc., version 9.1.2.226, San Diego, CA, USA) and the half maximal effective concentration (EC_50_), expressed as the relative enrichment factor of the extract in the bioassays, was determined by non-linear regression with a built-in log-logistic model. Toxicity was expressed in toxic units (TU) and was calculated by 1/EC_50_.

**Table 1.** Overview of bioassay battery applied to assess toxicity at four sites in Curaçao. TU = toxic unit, …EQ/L = equivalent concentration of the reference compound, n/a = not available. ^a^ = Van der Oost et al. (2017), ^b^ = Escher et al. 2018, ^c^ = Brion et al. (2019), ^d^ = this study.

|  | Bioassay | Endpoint | Reference compound | EBT | iEBT<sup>d</sup> | Unit |
| --- | --- | --- | --- | --- | --- | --- |
| *in vivo* | Bacterial bioluminescence inhibition | Luminescence inhibition | n/a | 0.05<sup>a</sup> | 0.005 | TU |
|  | Algal | Photosynthetic inhibition | n/a | 0.05<sup>a</sup> | 0.005 | TU |
| *in vitro* | PAH | Xenobiotic metabolism | benzo(a)pyrene | 6.21<sup>b</sup> | 0.621 | ng BEQ/L |
| | ERα | Estrogenic activity | 17β-estradiol | 0.28<sup>c</sup> | 0.028 | ng EEQ/L |

**Table 2.**
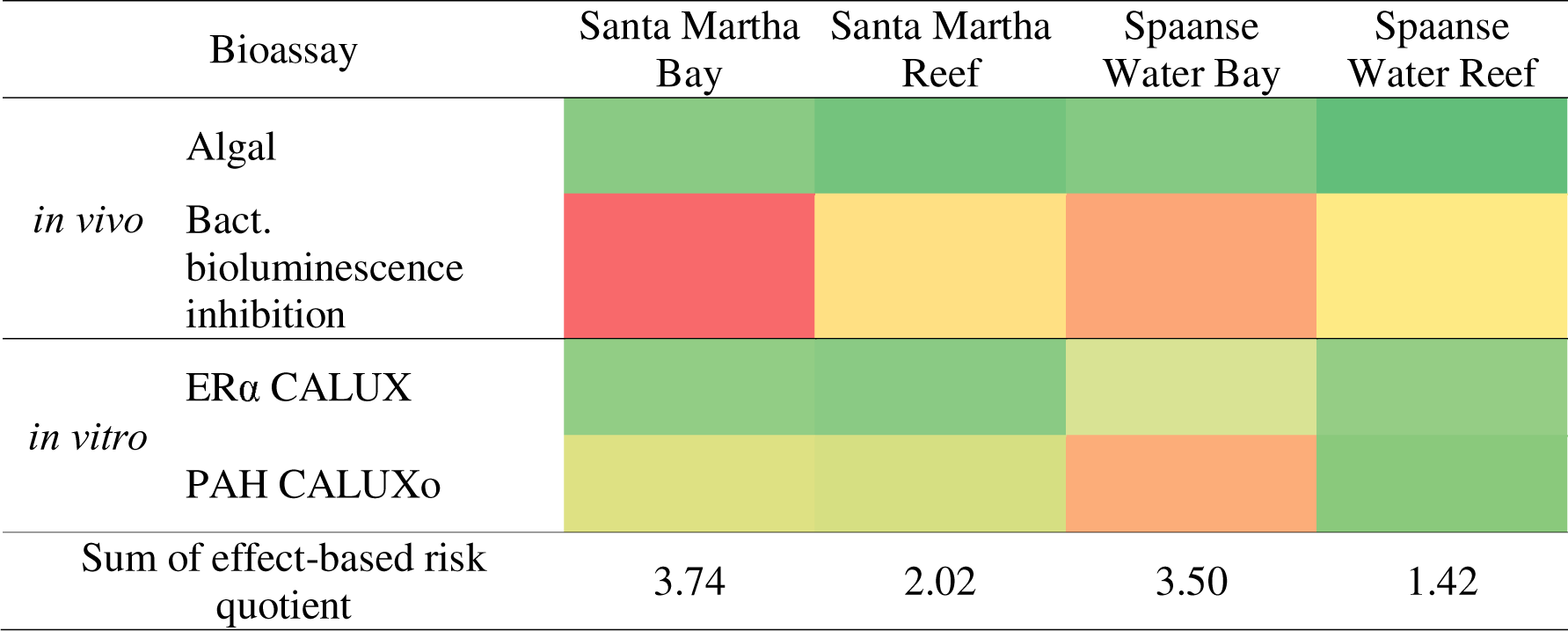
Heat map depicting the responses of four bioassays and the sum of the effect-based risk quotients for four sites during the wet season (Oct/Nov 2020). The colour gradient reflects the bioassay response value, in which red indicates the highest value (i.e. higher pollution levels) and green the lowest value (i.e. lower pollution levels) for that specific bioassay.

Two *in vitro* chemically activated luciferase gene expression (CALUX) reporter gene assays were applied to assess the activation of the estrogen and aryl hydrocarbon receptors (ERα and AhR), respectively. The two CALUX assays represent specific toxic endpoints and can serve as proxies for specific sources of pollution (Burg et al., 2013; De Baat et al., 2020). Estrogen receptor agonism (ERα CALUX), indicative of endocrine disruption, was used as a proxy for wastewater pollution (Välitalo et al., 2016) and aryl hydrocarbon receptor agonism (PAH CALUX), indicative of the induction of xenobiotic metabolism, as a proxy for industrial pollution (Neff et al., 2005). The ERα and PAH CALUX assays were performed by the BioDetection Systems laboratory (Amsterdam, The Netherlands) as described by De Baat et al., (2020). The effects of the extracts in the assays were expressed as bioanalytical equivalent (…EQ/L) of the reference compounds of the assays using the estimated sampled water volume of the passive samplers (Table 1). Further details on the bioassays are provided in the Supplemental Information.

#### Effect-based trigger values (EBT)

Effect-based trigger values (EBTs) were used to identify potential ecotoxicological risks based on the bioassay responses (Neale et al., 2023). The EBTs available in the scientific literature are commonly derived for risk assessments in freshwater systems. The presently used EBTs were derived by Brion et al., (2019) (ERα), Escher et al., (2018) (PAH) and van der Oost et al., (2017) (algal and bacterial inhibition) (Table 1). To render these EBTs suitable for the present marine risk assessment, an additional safety factor of 10 was applied to account for the generally higher sensitivity of marine organisms to toxicants (RIVM, 2015), resulting in interim effect-based trigger values (iEBT) (Table 1). To calculate ecotoxicological risk quotients, the bioassay responses were divided by their respective EBTs or iEBTs, where an effect-based risk quotient ≥ 1 indicated a potential ecotoxicological risk (De Baat et al., 2020). Cumulative ecotoxicological risk for each site was subsequently obtained by summing the four effect-based risk quotients for each site. This sum allowed the ranking of sites based on their ecotoxicological risk, where the highest sum of a site is assumed to represent the highest chemical burden, resulting from organic chemical pollution.

### Statistical analysis

Values in the raw dataset were examined for the presence of extreme outliers and excluded when they exceeded 3x the interquartile range (IQR). The effect of habitat (bay vs reef) and season (dry vs wet) on the daily average and the daily variability of all environmental parameters (i.e. temperature, pH, DO, salinity and PAR) was statistically analysed using the nonparametric Scheirer-Ray-Hare extension of the Kruskal-Wallis test with a post-hoc Dunn test because the assumptions of normality were violated (Scheirer et al., 1976). The assumptions of a normal distribution and equal variance were tested with a Shapiro-Wilks test and F-test, respectively. The Scheirer-Ray-Hare extension with a post-hoc Dunn test was also used to test for differences in average nutrient concentrations in the water samples between habitat and season since the assumptions of normality and homogeneity of variance were not met. This also applied to the total alkalinity and carbonate chemistry data. All statistical analyses were performed using Rstudio software (Version 2022.07.2+576 "Spotted Wakerobin"). P-values ≤0.05 were considered significant.

## Results

### Benthic community assessment

Benthic cover surrounding the monitoring stations in Spaanse Water Bay was dominated by seagrass and macroalgal meadows (48% ± 7 SE and 23% ± 3 SE, respectively), while all other monitoring locations were dominated by rubble and soft sediment (Figure 2). Spaanse Water Bay was characterized by the presence of the benthic upside-down jellyfish *Cassiopea* sp. (0.08%), whereas Santa Martha Bay had a high prevalence of the echinoid *Diadema* sp. (0.98%). Benthic community surveys revealed a strong difference in coral cover between inland bays (< 1%) and reefs (∼ 8%) (Figure 2). Specifically, coral cover (all species included) in Spaanse Water Bay was 12 times lower than in Spaanse Water Reef and 5.5 times lower in Santa Martha Bay than in Santa Martha Reef. Similarly, coral diversity was substantially lower in the bays than on reef flats. In total, four coral species were identified in the monitoring area at both bays, whereas 15 species (including one unknown) were identified in the monitoring area on the reefs (Table S5).

**Figure 2.**
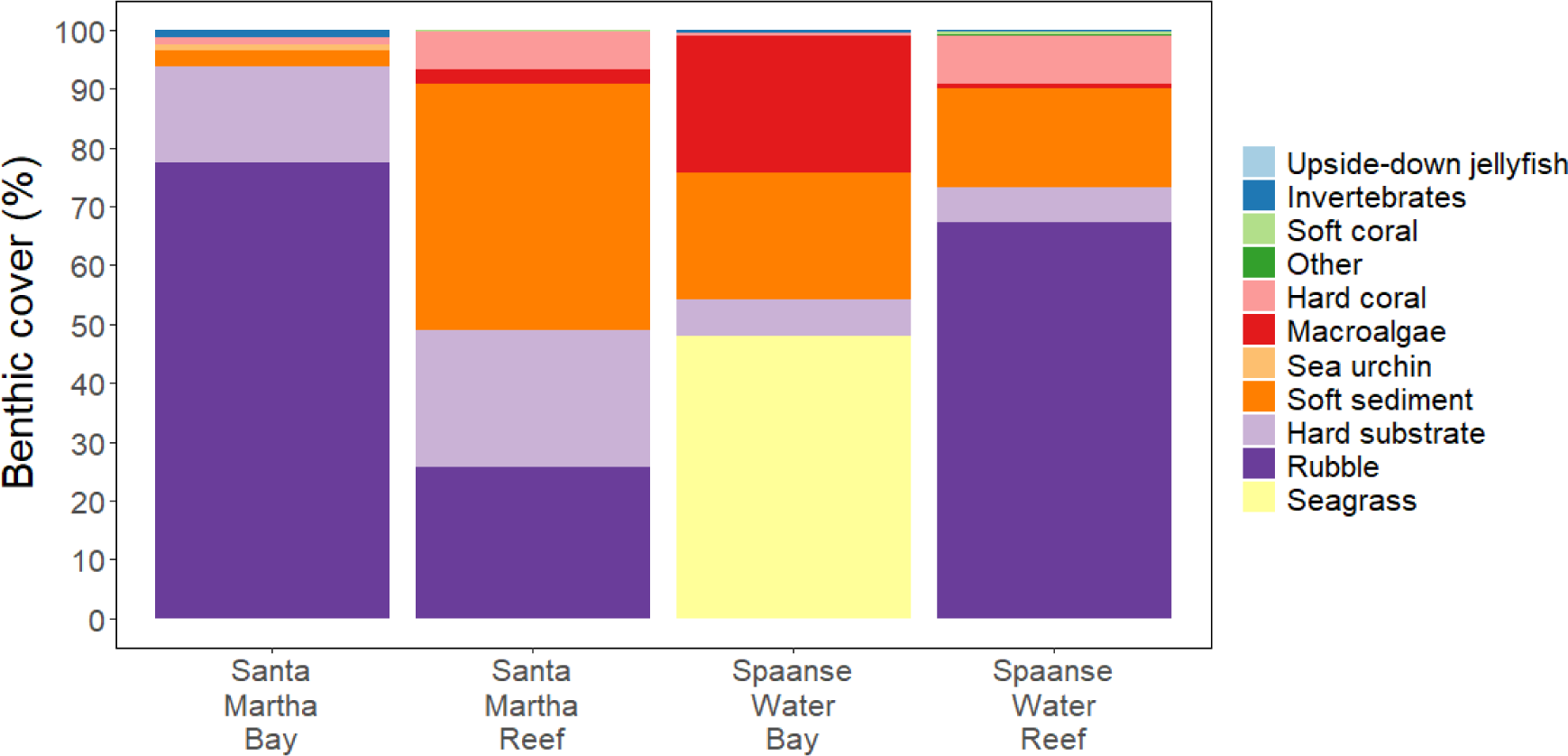
The benthic community at the four sites, expressed as cover percentage (%). The ‘invertebrates’ category comprises anemones, sponges, crustaceans (e.g. *Panulirus argus* and *Mithrax spinosissimus*) and feather duster worms. The ‘other’ category includes benthic cyanobacterial mats (BCM) and dead coral.

### Temperature, salinity, pH_T_, DO and PAR

The assessment of physicochemical conditions revealed that the bays not only had significantly different average conditions but were also much more variable than the reefs (Figure 3, Tables S6-8). Furthermore, strong differences between the dry and wet seasons were also observed for most parameters, while some showed complex interactive effects.

**Figure 3.**
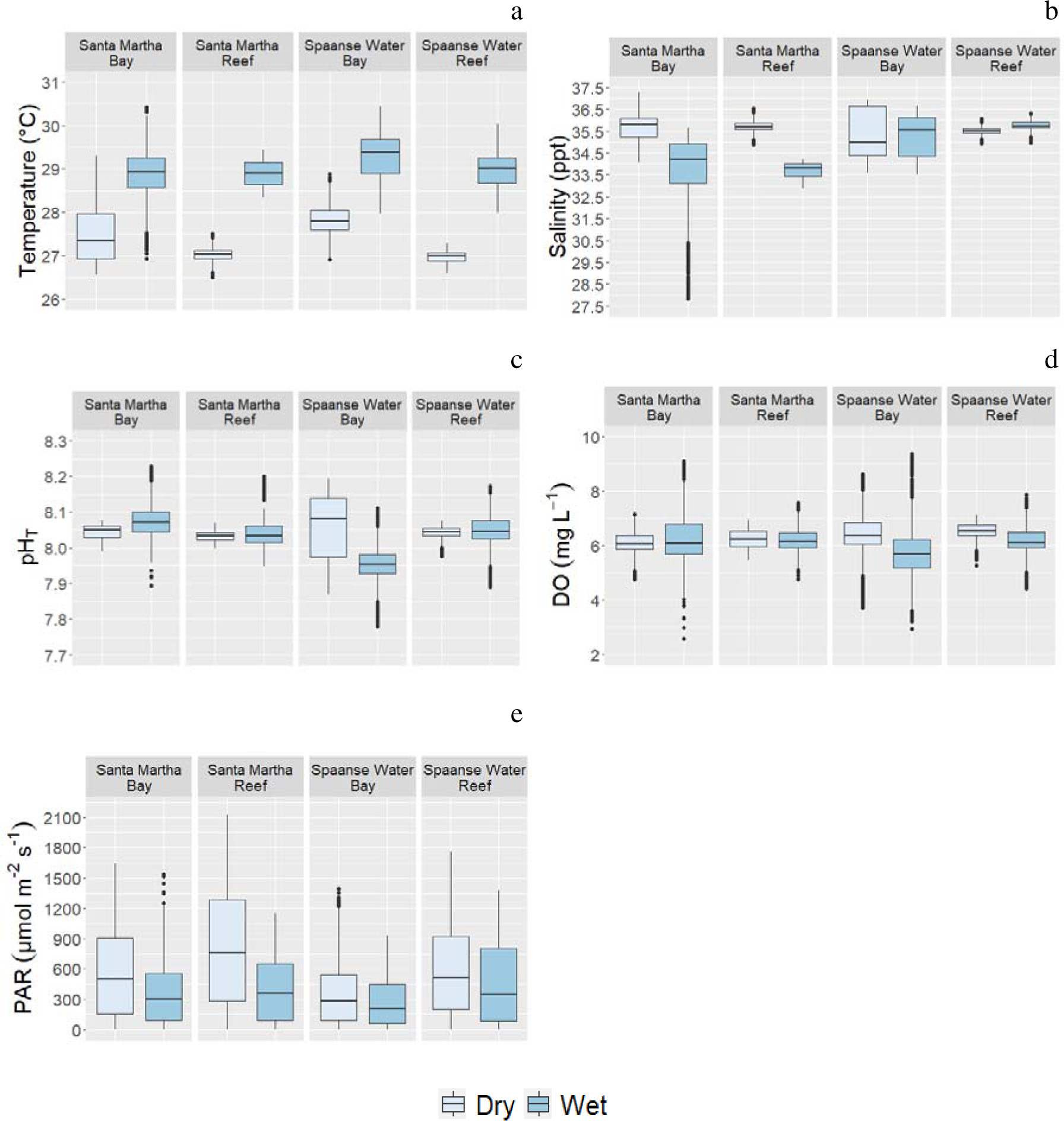
Physicochemical conditions measured in March 2020 (cool, dry season) and Oct/Nov 2020 (warm, wet season). a. temperature (°C), b. salinity (ppt), c. pH_T_ d. dissolved oxygen (mg L^-1^), e. photosynthetically active radiation (µmol m^-2^ s-^1^). Boxes span the first to third quartiles; the horizontal line inside the boxes represents the median, and black dots represent the outliers between the range of 3 times the IQR as extreme outliers were excluded that exceeded 3 times the IQR. Whiskers represent the range of the data that falls within 1.5 times the IQR. See also Table S1.

The average temperature in the bays was significantly higher (up to 0.8°C) than on the reefs (p = 0.043) (Figure 3a, Tables S6, S7). The daily temperature variability was significantly higher in the bays (up to 2.6 °C in the dry season and up to 2 °C in the wet season) compared to only up to 0.8 °C in the dry and wet season on the reefs (p = ≤ 0.001) (Figure 3a, Tables S6, S8). In addition, temperature was more variable in the dry compared to the wet season (p = 0.003) (figure 3a, Tables S6, S8). Notably, maximum temperatures exceeded the local coral bleaching threshold of 29.0°C (MMM = 28 °C + 1 °C; Coral Reef Watch, virtual station Aruba, Curaçao, and Bonaire, NOAA 2020-2021) by up to 1.4°C in the bays during the warmer dry season, while this was not the case for the reefs (Table S6). Average temperatures were generally warmer during the wet than the dry season but daily temperature variability did not differ significantly between the seasons (p = 0.074) (Figure 3, Tables S6-S8).

The average salinity was not significantly different between bays and reefs (p = 0.680), however, there was a significant difference between seasons (p = 0.005), with higher values in the dry (by 0.85 ppt) compared to the wet season (Figure 3b, Tables S6, S7). Daily variability of salinity differed significantly between both habitats (p ≤ 0.001) and seasons (p = 0.002), but no significant interactive effect was found (p = 0.944). Salinity was generally more variable in the bays than on reefs (up to 7.8 ppt versus up to 1.7 ppt daily), and also during the wet compared to the dry season (up to 3.4 ppt versus up to 2.4 ppt daily) (Figure 3b, Tables S6, S8).

The reefs had a significantly higher average pH_T_ than the bays (p = 0.033), although most sites had a relatively similar average pH_T_ within 8.03–8.08 during both seasons (Figure 3, Tables S6, S7). The only exception was Spaanse Water Bay during the wet season which had the lowest average pH_T_ found (7.95) (Figure 3c, Table S6). Average pH_T_ did not differ significantly between the dry and wet seasons (Figure 3c, Tables S6, S8). In contrast, the daily pH_T_ variability tended to be higher in the bays than the reefs, but only during the wet season (p = 0.03), and variability was generally higher during the wet compared to the dry season (Figure 3c, Tables S6, S8).

The average dissolved oxygen (DO) concentration was significantly higher (2.2%) on the reefs than in bays (p = 0.002), and also 3.0% higher during the dry than wet season (p ≤ 0.001) (Figure 3d, Tables S6, S7). The daily DO variability differed significantly between habitats (p = ≤ 0.001) and between seasons (p = ≤ 0.001), with 56.6% higher daily DO variability in the bays than on the reefs and during the wet season compared to the dry season. A significant interactive effect between habitat and season was observed for the daily variability of DO (p = 0.028) but not for average DO concentration (Figure 3d, Tables S6-S8). During the dry season, Spaanse Water Bay reached dissolved oxygen levels as low as 3.72 mg L^-1^, while Santa Martha Bay reached DO levels as low as 2.58 mg L^-1^ during the wet season.

Both average photosynthetically active radiation (PAR) and daily variability in PAR differed significantly between habitat and season (all p ≤ 0.001), however, no interactive effect was found between habitat and season (p = 0.957; p = 0.661) (Figure 3e, Tables S6-S8). The bays experienced 31.5 % lower PAR than the reefs, and PAR was 33% lower during the wet than the dry season. The maximum PAR was up to ∼2100 µmol m^-2^ s^-1^ on the reefs but only up to ∼1600 µmol m^-2^ s^-1^ in the bays (Figure 3e, Table S6).

### Seawater carbonate chemistry

These results are described in more detail in the Supplementary Materials (Figure S5, Table S12).

### Seawater nutrient concentrations

Overall, nutrient levels were significantly higher during the wet than the dry season and the bays tended to have higher nutrient levels than the reefs, particularly during the wet season (Figure 4, Table S9). Nevertheless, some of these effects differed for the different types of nutrients. Both nitrate and phosphate concentrations were 81% and 24% higher during the wet than the dry season, respectively, but did not differ significantly between habitats (Figure 4a, b; Table S9). Similarly, ammonium concentrations were also 68% higher during the wet than the dry season but were the only nutrient type that differed significantly between habitats, with the bays having 13% higher ammonium concentrations than the reefs (Figure 4c; Table S9).

**Figure 4.**
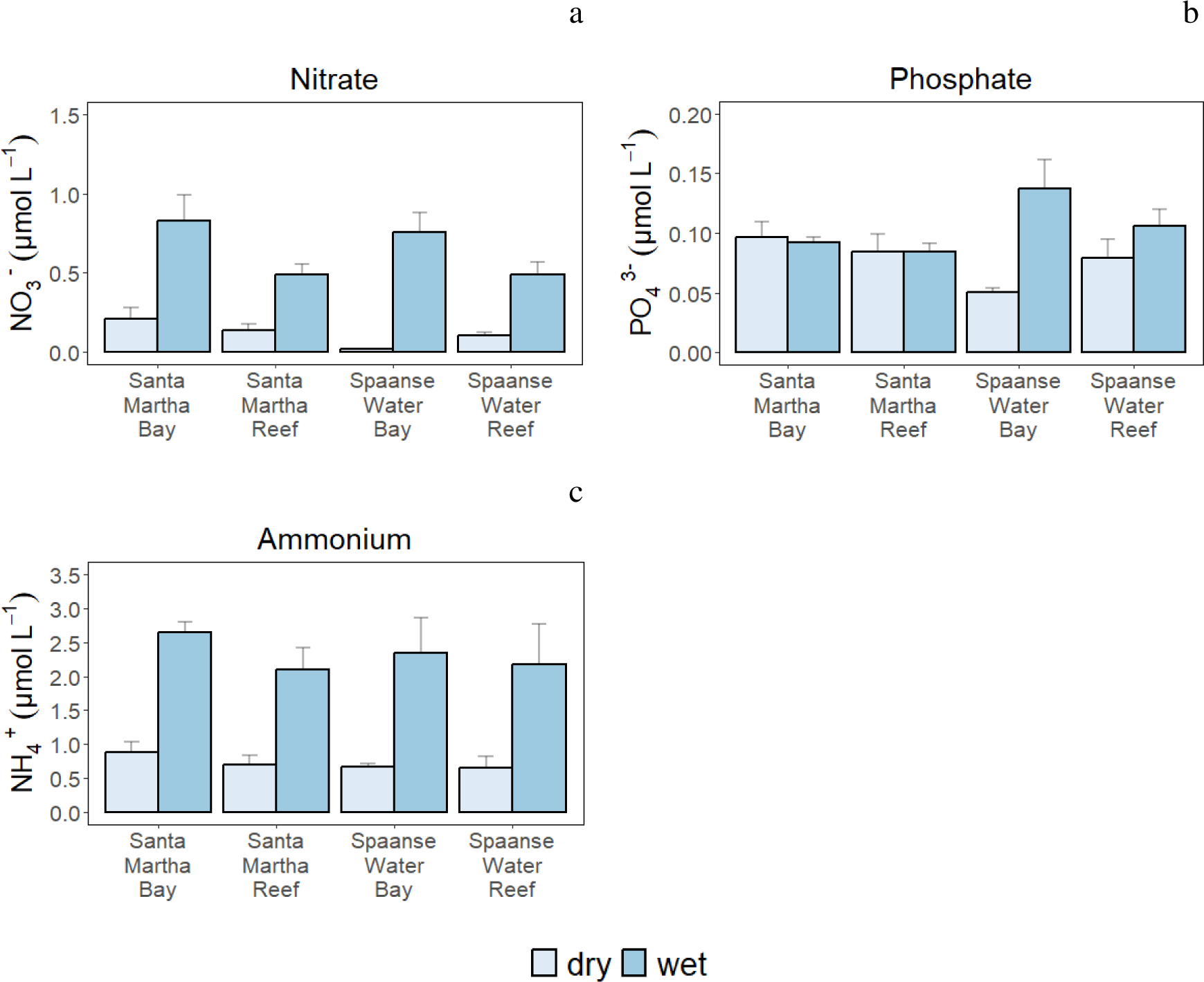
Nutrient concentrations measured in March 2020 (cool, dry season) and Oct/Nov 2020 (warm, wet season). a. Nitrate (NO_3_^-^), b. Phosphate (PO_4_^3-^) c. Ammonium (NH ^+^). Bars represent means ± the standard error (SE).

### Seagrasses as indicators of nutrient and trace metal pollution

In contrast to the water samples, the seagrass samples were only used to examine bay-specific nutrient and trace metal pollution as seagrass was only found in the bays. These results are described in more detail in the Supplementary Materials (Tables S10 and S11).

### Sediment trap collection rates

Average trap collection rates during the wet season ranged from 0.94 to 3.23 mg cm^-2^ day^-1^, whereby Santa Martha Reef had the highest rate (3.23 mg cm^-2^ day^-1^), and Spaanse Water Bay the lowest (0.94 mg cm^-2^ day^-1^) (Figure 5a). The reefs received overall higher sediment loads than the bays, however, the sediment particles in the bays had a higher organic matter content (∼11.5% versus ∼4.5% on the reefs) (Figure 5b). Santa Martha Bay sediment particles had the highest organic matter content (0.29 mg cm^-2^ day^-1^), which was ∼ 50% higher compared to the other three sites. The traps collected a high percentage of the smaller particles as a percentage of the total weight, whereas the opposite was true for the sediment collected from the benthos (Figure S3). Furthermore, trapped sediment on the reefs was dominated by smaller sizes compared to the bays. The benthic sediment showed the opposite, with bays having a higher percentage of large particles than the reefs, particularly Santa Martha Bay.

**Figure 5.**
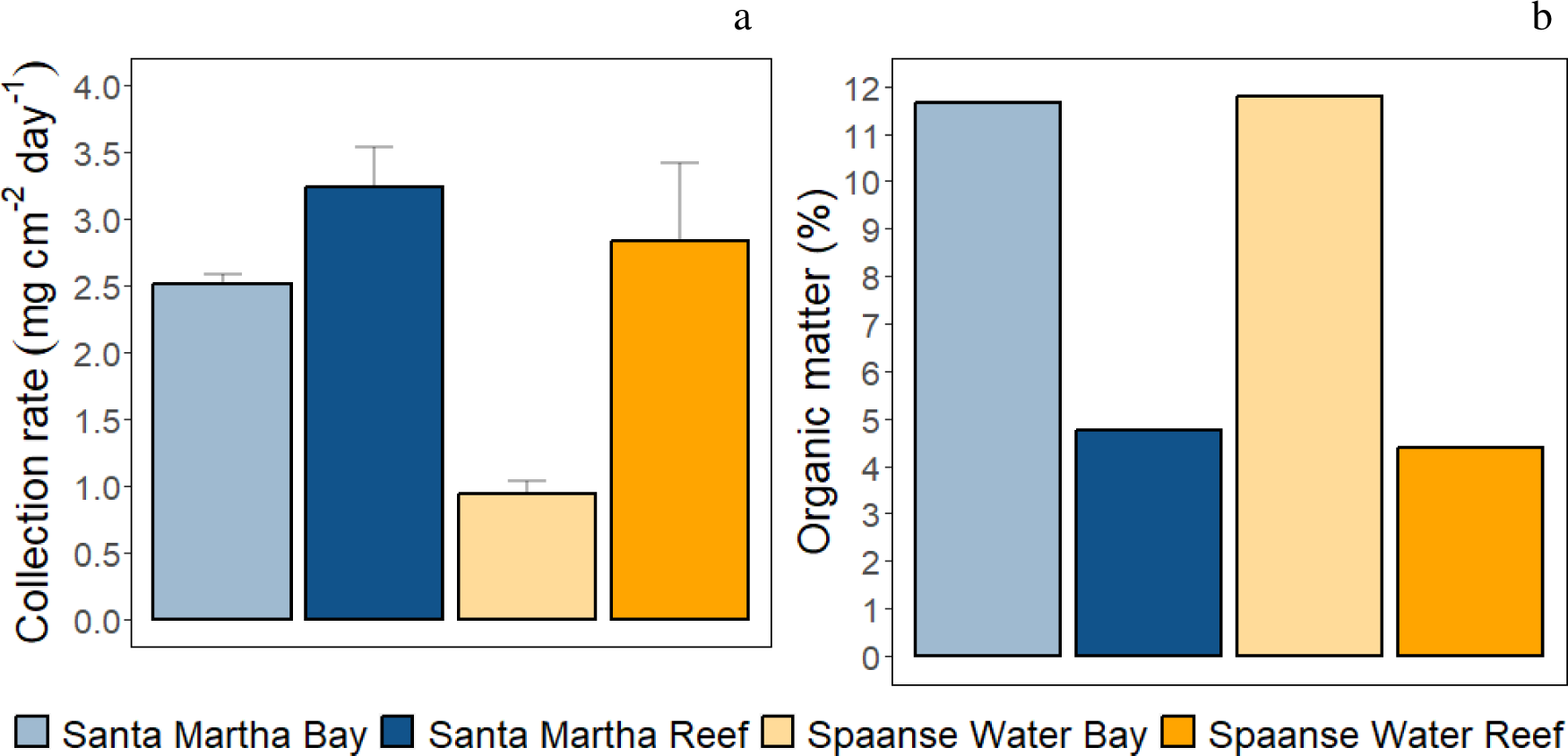
a. Average trap collection rates (mg cm^-2^ day^-1^) for all four sites during the warm wet season (Oct-Nov 2020), Shown is mean ± SD. b. Organic matter percentage of the total sediment weight accumulated in the sediment trap.

**Figure 6.**
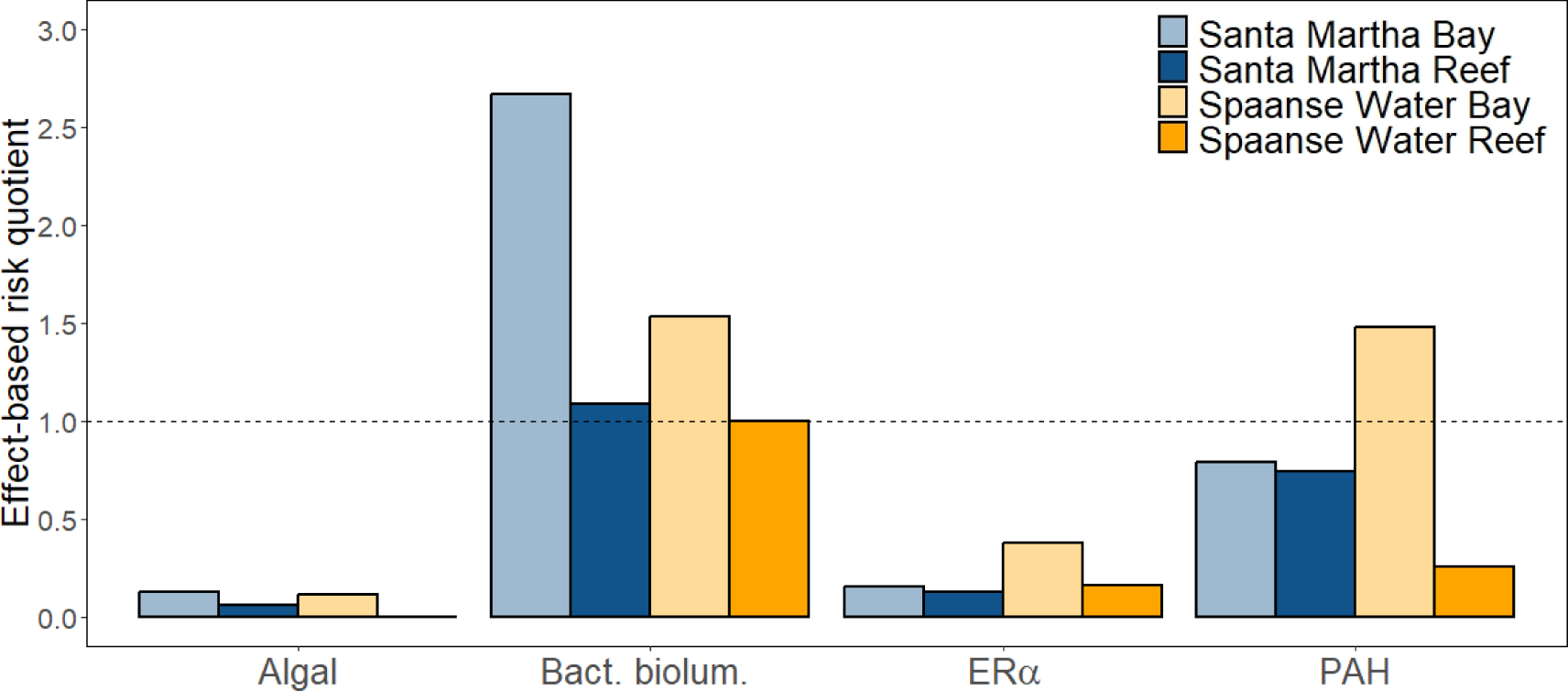
Responses of four bioassays at four sites during the wet season (Oct/Nov 2020). The dotted line represents an effect-based risk quotient ≥1, derived by dividing the bioassay response by the intermediate effect-based trigger value for the marine environment (iEBT). Exceedance of this value indicates a potential ecotoxicological risk. Algal = Algal bioassay, Bact. biolum. = Bacterial bioluminescence inhibition assay, ERα = Estrogen receptor α CALUX (a proxy for wastewater), PAH = polycyclic aromatic hydrocarbon CALUX (a proxy for industrial pollution).

### Effect-based chemical water quality assessment

All bioassays met their respective validity criteria and responses were observed in all bioassays. While the EBT was exceeded at none of the sites (Figure S4), the more conservative iEBT was exceeded (Figure 4). For the algal bioassay, responses above the iEBT threshold value were not detected but the passive sampler extracts from the bays caused higher responses than those from the reefs, and Spaanse Water Reef extracts caused no bioanalytical response at all. In the bacterial bioluminescence inhibition assay, the iEBT threshold value was exceeded at Santa Martha Bay, Santa Martha Reef and Spaanse Water Bay, while the response at Spaanse Water Reef was on par with the iEBT. Again, the highest responses for this assay were detected at the bay sites. The ERα CALUX assay, a proxy for wastewater, showed the highest response in Spaanse Water Bay but the iEBT was exceeded at none of the sites. The PAH CALUX assay revealed that industrial pollution was present at each site, but only Spaanse Water Bay exceeded the iEBT. The bays showed usually higher responses in comparison to the reefs, which is also reflected in the exceedance of the iEBT in the bioassays (Figure 4).

A summary of the bioassays battery responses is shown in a heat map (Table 7). Spaanse Water Bay extracts caused the highest responses in all bioassays, followed by Santa Martha Bay, Santa Martha Reef and Spaanse Water Reef. Hence, the bays exhibited a higher cumulative mixture toxic pressure than the reefs.

## Discussion

### Are Curaçao’s inland bays potential analogues for future ocean conditions?

The presently studied inland bays represent thermally extreme environments because they were overall up to 0.8°C warmer than the reefs and had high daily temperature variability year-round (up to 2.6°C) (Fig. 3a, Table S6). This makes them somewhat comparable to, though less extreme than, other semi-enclosed lagoons, such as the Bouraké lagoon in New Caledonia (up to 6.5°C, Maggioni et al., 2021) or the mangrove lagoons in the Great Barrier Reef with a temperature range up to 7.7°C across three months compared to 3.5°C in this study (Camp et al., 2019). However, when compared to a mangrove site Belize, where the average daily temperature variance was 0.58°C (Lord et al., 2021), the inland bays can be considered a more extreme site in the Caribbean Sea. Importantly, temperatures in the bays already exceeded the full annual monthly temperature range for the reefs (25.7°C – 28.0°C; Coral Reef Watch, virtual station Aruba, Curaçao, and Bonaire, NOAA, 2021) during the cool dry season. Furthermore, maximum inland bay temperatures during the warm, wet season were in line with the predicted mid-century sea surface temperature (global open ocean) under the intermediate emission scenario SSP2-4.5 (IPCC, 2019, 2021).

While mangroves, seagrasses and the organisms associated with these habitats are typically adapted to such dynamic temperature regimes, they represent extreme and potentially stressful temperatures for corals that have much lower thermal tolerance. Specifically, the local coral bleaching threshold is only ∼29°C (NOAA, 2021) whereas upper thermal limits of tropical seagrass species range from 32-38°C (Marbà et al., 2022). Thus, inland bay corals were exposed to temperatures exceeding their bleaching thresholds already during the cool dry season and spent most of their time at or above the bleaching threshold temperature during the warm wet season (Table S6). It should be noted here that temperatures during the 2020 wet season were warmer than usual (NOAA, 2021), but while some corals on the reefs were bleached at greater depth (10-15 m), this was not observed in the inland bays (although some corals appeared pale; CDJ, personal observation). These findings suggest that inland bay corals have greater heat tolerance than reef corals which is likely due to regular exposure to more variable temperatures (Palumbi et al., 2014; Rivest et al., 2017; Safaie et al., 2018; Schoepf et al., 2015).

The inland bays showed much larger daily fluctuations in DO compared to nearby reefs in both seasons, whereas for pH_T_ this was the case only in the wet season (Figure 3c, d, Table S6). This strong daily variability is driven largely by the high cover of seagrass, macroalgae, and mangroves in the bays (Figure 2), combined with long seawater residence times due to restricted exchange with the open ocean. These benthic communities influence the pH_T_ and DO by performing photosynthesis by day and respiration by night (Delille et al., 2000; Hendriks et al., 2014). Although corals also photosynthesize, the daily oscillations in pH_T_ and DO are typically much smaller in a coral-dominated habitat as the increase of pH_T_ and DO by photosynthesis of endosymbionts is counteracted by calcification and respiration of associated coral reef organisms (Anthony et al., 2013; Cryer et al., 2023; Page et al., 2016). The observed daily pH variability of up to 0.25 units is lower than the 0.69 daily range in the semi-enclosed lagoon system in Bouraké, New Caledonia (Maggioni et al., 2021). The pH range of 0.34 units found during the monitoring period in the inland bays was higher than the pH range of 0.21 units found in a mangrove forest in Panama (0.21 units) (Stewart et al., 2022). Similarly, the observed daily DO variability of up to 5.91 mg L^-1^ is comparable to 4.91 mg L^-1^ in the semi-enclosed lagoon system in Bouraké, New Caledonia (Maggioni et al., 2021), but the average DO found in the inland bays (6.1 mg L^-1^) is higher than the average DO (3.74 mg L^-1^) found in a mangrove forest in Panama (Stewart et al., 2022). The persistence of corals in such extreme environments is remarkable given their high sensitivity to low pH (Albright & Langdon, 2011; Kornder et al., 2018) and to some degree low DO (Altieri et al., 2017; Johnson et al., 2021). The inland bays are, therefore, ideal locations to investigate how such strong diel variability modulates resistance to acidification and severe de-oxygenation or hypoxia (Altieri et al., 2021; Rivest et al., 2017).

While average pH and DO levels in the inland bays were largely within the range of present-day conditions, the strong diel fluctuations of DO and pH led to regular but temporary exposure to future ocean conditions for resident organisms (Table S6). The most extreme site was Spaanse Water Bay where even average pH_T_ (7.95) was substantially reduced during the warm wet season, reaching levels predicted by ∼2050 under the high-emission SSP3-7.0 scenario (IPCC, 2021). Minimum pH_T_ values ranged from 7.78 to 7.87, which compares to predicted end-of-century pH levels of 7.9 and 7.75 under the SSP3-7.0 and SSP2-4.5 scenarios, respectively (IPCC, 2021). Similarly, the low average DO recorded in Spaanse Water Bay during the wet season (5.75 mg L^-1^) represented a 10% decrease compared to the dry season, which is more than the predicted global oxygen loss of ∼6% by 2100 under SSP3.-7.0 (IPCC, 2021). Notably, minimum DO levels recorded in the bays during the wet season were <3 mg L^-1^ and thus occasionally came close to hypoxic levels (<2 mg L^-1^). While the temporary exposure associated with strong diel fluctuations is vastly different from an acute hypoxia event (Lucey et al., 2020), these results nevertheless show a high risk for sustained hypoxia events to occur inside the bays during the wet season – indeed, such events may have already occurred but gone undocumented (Altieri et al., 2017). However, the strong diel DO and pH variability could also enhance the resistance of resident organisms to sustained hypoxia and acidification, as has been documented, for example, in corals for heat and to some degree acidification resistance (Brown et al., 2022; Kapsenberg & Cyronak, 2019; Rivest et al., 2017; Safaie et al., 2018).

Taken together, our results show that the inland bays of Curaçao have significant potential to serve as natural laboratories to study the effects of predicted future ocean conditions on resident taxa *in situ*. This applies particularly within the context of strong diel fluctuations rather than shifted mean conditions and with the caveat of co-occurring stressors. While a diverse range of natural laboratories has been documented in recent years (Burt et al., 2020; Camp et al., 2018), true natural analogues that perfectly simulate future ocean conditions rarely exist since co-occurring stressors are present at most of these locations. For example, the semi-enclosed lagoon of Bouraké, New Caledonia, has among the most extreme temperature, pH and DO conditions documented for a coral reef environment but also has lower light and higher levels of organic matter and certain nutrients than reference reefs (Maggioni et al., 2021). In Curaçao’s inland bays, where coral cover and diversity are much lower than in Bouraké (Figures 2, S5, Vermeij et al., 2007), co-occurring stressors include both natural and anthropogenic factors, since naturally high turbidity and sedimentation rates have been exacerbated by eutrophication and, in some areas, other forms of pollution (see next section). As human impacts on tropical marine ecosystems continue to increase, the presence of these stressors in the inland bays can offer a more realistic outlook into how global and local stressors will interact in the future but comprehensive, high-resolution monitoring is essential. Overall, Curaçao’s inland bays are unique natural laboratories to investigate how environmental variability modulates stress tolerance to future ocean conditions and could play an important role as resilience hotspots or refugia adaptation (Kapsenberg & Cyronak, 2019; Rivest et al., 2017; Schoepf et al., 2023).

### Are Curaçao’s inland bays threatened by local stressors?

The inland bays generally showed elevated nutrient levels compared to the reefs, particularly during the wet season, although this also depended on methodology (Fig. 2, Tables S4, S5). Given the lack of major rivers in Curaçao, nutrient input into the marine environment typically occurs in the form of rainfall-associated pulses during the wet season (Den Haan et al., 2016) as well as chronic seepage from septic tanks (Estep et al., 2017). Thus, time-integrated monitoring via bio-indicators provides more reliable estimates of nutrient levels than discrete water samples. Indeed, our seagrass samples indicated that both bays should be considered eutrophic coral habitats (Govers et al., 2014a; van Tussenbroek et al., 2016), with Spaanse Water Bay being slightly more eutrophied than Santa Martha Bay due to high coastal development. In contrast, water samples showed nutrient concentrations within the oligotrophic range (Crossland & Barnes, 1983; Tanaka et al., 2007) but are less representative because they only present a snapshot in time. The water samples, however, confirmed significantly higher nitrate, phosphate and ammonium levels during the wet than dry season across the mangrove-seagrass-coral reef continuum, but only ammonium was 13.4% higher in the bays than on reefs. Future research should consider the use of macroalgal bioindicators (Vaughan et al., 2021) because, in contrast to seagrass, macroalgae are present in all habitats along this continuum (Figure 2). Overall, elevated nutrient concentrations can have a wide range of effects on marine taxa (Nalley et al., 2023) and may make them more susceptible to co-occurring stressors such as coastal acidification (Silbiger et al., 2018), but this may not be the case for moderate nutrient concentrations in corals (Dobson et al., 2021).

The high turbidity of the inland bays also has natural causes as low visibility and high (inorganic) sediment trap collection rates were already reported ∼60-100 years ago (de Kock & de Wilde, 1964; van der Horst, 1924). Interestingly, in the present study, the two reef sites had higher sediment trap collection rates than the two bays (Figure 5a) but this could represent a potential influx from the bays that was enhanced during the wet season. In addition, the very different flow regimes make it challenging to compare trap collection rates between sheltered bays and exposed reefs because there is no apparent relationship between trap collection rate and turbidity in low-flow areas (Storlazzi et al., 2011). Comparison with previous work (e.g. Kuenen & Debrot, 1995) is also difficult because careful calibration experiments would be required to account for differences in trap design, deployment parameters and locations (Storlazzi et al., 2011). While high sedimentation rates can generally have negative impacts on the health of coral and other marine organisms (Fabricius, 2005; Rogers, 1990), this is particularly the case when organic matter content is high (Bartley et al., 2014; Weber et al., 2006). The inland bay sediment had more than twice as much organic content than reef sediment, especially in Santa Martha Bay, which can influence the water clarity, as finer particles settle more slowly, causing reduced light availability for longer periods, influencing the photosynthetic capability of organisms and consequently the local pH and oxygen conditions (De Boer, 2007; Fabricius et al., 2016; Storlazzi et al., 2015). It is therefore likely that the sediment regimes in the bays contribute to a more stressful environment for sensitive organisms such as corals (Bainbridge et al., 2018).

Seagrass leaves collected in the two inland bays revealed the presence of certain trace metals (Table S11) but it is unclear whether levels were higher than on the reefs given that no seagrass could be collected there. *Halophila stipulacea* - a bioindicator for Cu and Zn (Bonanno & Raccuia, 2018) - had much higher concentrations of both trace metals in Spaanse Water than in Santa Martha Bay. Given that Cu is used as a biocide in antifouling paints while Zn is used to create metal alloys and pigments, and as a fungicide (Nalley et al., 2021), this could well be related to the greater degree of coastal development and boating activities at Spaanse Water Bay (Govers et al., 2014b). In addition, *Thalassia testidinum* (collected in Spaanse Water Bay only) had Cu and Ni levels that were 20% and 69% higher, respectively than in the study by Govers et al., (2014b), whereas Fe and Zn concentrations were ∼50% lower. This suggests that trace metal concentrations in Spaanse Water Bay may have changed over time but that overall trace metal pollution is ongoing. Notably, it is important to acknowledge that Ni and Fe are naturally occurring elements in the marine environment. However, Ni can also be present in higher concentrations as a result of industrial pollution, whereas Fe is not only essential for plants and animals but is also widely utilized in various manufacturing processes (Nalley et al., 2021). Several other trace metals were not detected, such as As, Cd, Co, Pb and Se, but should be considered when determining the health of resident marine organisms as stress could be intensified by the complex interaction between trace metals and global stressors (Kibria et al., 2021; Negri & Hoogenboom, 2011).

### Effect-based chemical water quality assessment

The combined use of passive sampling and effect-based methods represents a promising approach to study chemical water quality (De Baat et al., 2020), but has so far seen limited application in tropical reef ecosystems (Shaw et al., 2009) highlighting the novelty of the present study. After the comparison of the bioassay responses to the respective threshold values, potential ecotoxicological risks were detected using the bacterial bioluminescence inhibition assay at all four sites. This assay is both indicative of general cytotoxicity in environmental samples as well as toxicity to marine bacteria. Hence, the detection of ecotoxicological risks using the bacterial bioluminescence inhibition assay is relevant as it indicates a concerning level of organic chemical pollution at all sites. Additionally, this is of concern since bacteria dominate the ocean in abundance and support the existence of higher life forms (Azam & Malfatti, 2007). Moreover, corals are holobionts, consisting of the coral itself and the associated algal symbionts, bacteria and, fungi (Cleary et al., 2019; Knowlton & Rohwer, 2003), and coral-associated microorganisms could be affected by the toxic compounds present in their environment. This suggests that coral health may be reduced because coral-associated bacteria have many beneficial roles for different coral life stages (McDevitt-Irwin et al., 2017). The interim effect-based trigger value of the algal photosynthetic inhibition assay, on the other hand, was not exceeded at any site, indicating that there appear to be low risks of chemicals inhibiting photosynthesis, such as herbicides (Marzonie et al., 2021).

The *in vitro* CALUX assays confirmed the presence of organic chemical pollution and pointed towards industrial- and wastewater effluents as sources of pollution. However, Spaanse Water Bay was the only site at which the interim effect-based trigger value for the proxy for industrial pollution (PAH CALUX) was exceeded. This is explainable since Spaanse Water Bay is near Caracas Bay, which was previously used as an oil storage facility (1920-1986) (Debrot et al., 1998). Over the years, petroleum spillage and leakage may have caused significant and persistent hydrocarbon pollution in which the sediment currently serves as a source of legacy industrial pollution to the overlying water. Especially in the more turbid wet season, resuspension of polluted sediments may have caused increased levels of hydrocarbons in the overlying water, causing exposure of organisms to toxic chemicals, reflected in the higher PAH CALUX response recorded at this location. Co-exposure to petroleum pollution and ultraviolet radiation poses significant risks to the early life stages of corals (Nordborg et al., 2021). The interim effect-based trigger value for the proxy for wastewater pollution (ERα CALUX) was not exceeded at any of the investigated sites, but the highest response for this bioassay was also found in Spaanse Water Bay. This could likely be due to the wastewater seepage that has been increasing since the bay shifted towards a residential character in the 1980s (Debrot et al., 1998; Kuenen & Debrot, 1995). This effect can be exacerbated by the wet season but apparently did not cause ecotoxicological risks at the present location and period of sampling.

The cumulative effect-based risk quotients obtained in the present study indicate that ecotoxicological risks are potentially present at all the investigated locations. This pervasive mixture-toxic pressure was effectively detected by combining passive sampling and a bioassay battery. This highlights the value of combining these techniques, and underlines the relevance of applying effect-based methods for chemical water quality assessment, also in tropical ecosystems (Caracciolo et al., 2023; De Baat et al., 2020). Even the modest set of two *in vivo* and two *in vitro* bioassays, representing two relevant reef organism groups and two relevant modes of action, elucidated the effects of a wide range of bioavailable contaminant mixtures in the environment. In future effect-based water quality assessments of coral reef environments, the set of applied bioassays can be expanded to include the oxidative stress response as well as bioassays targeting integral life stages or functions of coral reef biota such as scleractinian coral and sea urchin larvae (Neale et al., 2023; Shaw et al., 2009). The highest cumulative effect-based risk quotient was found for Spaanse Water Bay, followed by Santa Martha Bay, Santa Martha Reef and Spaanse Water Reef. This clearly illustrates that organisms in the two inland bays are exposed to higher toxic pressures and the resulting ecotoxicological risks than organisms inhabiting the reefs. This further confirms that inland bays can serve as sentinels for future ocean conditions, also for the increasing chemical burden that threatens ecosystems worldwide (Persson et al., 2022).

Chemical pollution can have both direct and indirect toxic effects on coral reef organisms, yet these detrimental effects on tropical reef ecosystems have received little attention, especially when compared with the effects of climate change like increasing sea surface temperature and ocean acidification (Ouédraogo et al., 2023; Sigmund et al., 2023). In a recent study by Nalley et al., (2021), a systematic review and meta-analysis provided water quality thresholds for corals. Similarly, a systematic review of experimental studies by Ouédraogo et al., (2023) evaluated the toxicity of chemical pollution to tropical reef-building corals, producing usable data for ecological risk assessment. However, these chemical guidelines do not account for all possible chemicals that may be present in water, nor do they consider the mixture effects that may arise from the numerous chemicals simultaneously present in the water. Effect-based methods account for these effects of mixtures of (un)known compounds, but their application in the mangrove-seagrass-coral reef continuum has, so far, been limited, and unlike for freshwater systems, no well-established effect-based trigger value (EBT) for coral reef systems is currently available. As a precautionary and pragmatic approach to incorporate the potential sensitivity of coral reef organisms to chemical pollution, an additional safety factor of 10 was applied to the freshwater EBTs of the bioassays used in the present study, to extrapolate freshwater to marine water quality criteria (RIVM, 2015). However, it is presently unknown if the intermediate EBTs are sufficiently protective for coral reef organisms, or whether they are too strongly on the side of caution. Hence, the development of EBTs tailored to tropical reef ecosystems is warranted to facilitate water quality monitoring and the protection of these vulnerable and valuable ecosystems from the harmful effects of chemical pollution.

## Conclusion

This study used a multi-disciplinary approach to assess both biotic (i.e., benthic cover and coral diversity) and more than 20 abiotic parameters characterizing two inland bays and two nearby reference reefs in the southern Caribbean during the cool, dry season and warm, wet season. Our multi-pronged approach combined high-resolution temporal monitoring of key environmental parameters with time-integrated pollution monitoring using both bio-indicators (nutrient and trace metal pollution) and passive samplers and bioassays (organic chemical pollution). The inland bays of Curaçao are marginal and/or extreme environments where the impacts of global stressors in combination with local stressors are present. These bays can therefore act as natural laboratories to examine the effects of a changing ocean on marine organisms, but a caveat may be the presence of local stressors. However, local stressors have become significant threats to marine ecosystems and as they will not be reduced in the near future, there is a need to understand the combined effects of global and local stressors on marine organisms, which can be studied in these inland bays. In consideration of the vital link between the mangrove-seagrass-coral reef continuum offered by the inland bays, it is important to note that possibly the resident organisms are already at their limits in terms of their capacity to persist in these multi-stressor habitats. It is therefore highly needed for future research to identify their physiological mechanisms. The complexity of a multi-stressor environment such as the inland bays offers great opportunities and requires prolonged monitoring, which can ultimately result in the protection of these habitats as the ecological value of the inland bays in terms of marine life (e.g. mangrove, seagrass beds and seagrasses) is high.

## Acknowledgments

We thank: the Carmabi Research Station in Curaçao; Mark Vermeij for his advice; Svenja Pfeifer for assistance in the field; Rene van der Zande for constructing sediment traps; Pieter Slot for analyzing nutrient samples; Merijn Schuurmans for assistance with field and lab equipment; Bas van Beusekom for assistance with field and lab equipment and for his guidance during the algal bioassay; Rebecca van Oostveen for analyzing the benthic survey data; Titus Rombouts for assistance with the laboratory work of the sediment samples; Rutger van Hall for analyzing the nutrient concentration and trace metal concentration of the seagrass leaves; Michel van Son for assistance with the titrations; Peter Cenijn for the performance of the bacterial bioluminescence inhibition assay.

