## Supplemental Information for "Sentinels for future coral reef conditions: assessment of environmental variability and water quality in semi-enclosed inland bays in the southern Caribbean"

2. Present address: Dept. of Zoology, Stockholm University, Svante Arrhenius Väg 18b, 11418, Stockholm

† Contributed equally.

### **Supplemental Material and Methods**

#### **­­Study sites**

Local anthropogenic stressors – Santa Martha Bay is located in the less densely populated western part of Curaçao and surrounded only by a small village and fishing harbour further into the bay. The channel connecting SMB to the open ocean has been opened and closed in the past and was last reopened in 1964 (de Kock & de Wilde 1964). Contrastingly, Spaanse Water Bay is strongly influenced by local stressors. Over the previous decades, intensive coastal development has shifted the bay from primarily recreational to largely residential (Debrot et al., 1998; Kuenen & Debrot 1995). However, public sewage infrastructure is inadequate, resulting in sewage seepage into the bay (Debrot et al., 1998), while the many boats using this bay as a harbour add to eutrophication. Therefore, the two inland bays differ in the degree of local anthropogenic stressors.

Monitoring areas - The selection of monitoring areas within the bays was based on various factors, including accessibility, relatively high abundance of corals, and presence of mangroves to assess how their metabolism alters local seawater chemistry. Since coral abundance is very patchy inside the bays and mangroves surround most of SWB but only certain coral-inhabited areas of SMB, the monitoring location inside SMB was located ca. 100 m to the west of the entrance channel at 1.5 m depth, whereas it was located at 2.5 m depth at the end of one of the “fingers” in the south-western part of SWB (Fig. 1B, C). For the nearby reef sites, the monitoring areas were on the reef flat at ~5 m depth as Vermeij et al., (2007) as this is where most of the coral species present in the bays were also abundant, thus allowing for a more standardized comparison between sites.

#### **Environmental monitoring of temperature, salinity, pH_T_, DO and PAR**

Not all parameters were logged continuously for the full extent of the study period at each site due to logger malfunctions and high calibration frequency needed for the pH loggers. Therefore, some loggers were shuffled among sites (PAR and conductivity & temperature), while others were retrieved every other day for calibrations (pH) (Table S1). Loggers were placed on a cement block, ensuring that the PAR sensor and conductivity cell were vertically mounted (Fig. S1). The cement blocks were deployed at a depth of 1.5 m for Santa Martha Bay, 2.5 m for Spaanse Water Bay, 5.5 m for Santa Martha Reef, and 5.0 m for Spaanse Water Reef.

Conductivity and temperature – To prevent interference by other loggers, the conductivity logger was placed 10 cm away from any other object on the cement block. Loggers were calibrated with the factory-provided calibration equations using the Odyssey data logging software. For temperature, a linear calibration was applied, whereas for conductivity a polynomial calibration was applied. Before deployment, the accuracy of the conductivity cells was examined with a sodium chloride solution of 35 ppt at room temperature. Conductivity (mS/cm) was converted to salinity (ppt) with the 'marelac' package in R (Soetaert et al., 2010). For this function, 3 parameters were needed, i.e., the conductivity ratio, temperature and pressure against the local atmospheric pressure. To obtain the conductivity ratio, conductivity was divided by a standard conductivity of seawater at salinity = 35 ppt, temperature = 15°C, and pressure = 0 bar. A temperature of 25 °C for all conductivity data was used for the conductivity/salinity conversion^[1]^. The pressure used for the conversion is referenced against the local atmospheric pressure in bar and calculated with the following formula:	$P= \rho*g*h$ where $P$ = pressure in Pascal (100.000 Pascal = 1 Bar), $\rho$ = density of seawater at a certain temperature, $g$ = gravity velocity (9.81 m/s^2^) and $h$ = depth in m. The average seawater temperature measured by the conductivity loggers during the monitoring period was used to determine$\rho$.

pH_T_ – To determine seawater pH on the total scale (pH_T_), loggers were calibrated every second-day using tris(hydroxymethyl) aminomethane (TRIS) buffer at two different temperatures as described in Dickson et al., (2007). The certified TRIS buffers were purchased from Prof. A. Dickson (Scripps Institute of Oceanography). It was logistically not possible to calibrate the pH loggers every day at all four sites and therefore, only data from 12 hours before and up to 24 hours after calibration were used. The loggers were deployed with anti-fouling guards, which were always removed before calibration. pH_T_ was calculated using the mV and temperature output of the pH loggers. First, a linear regression, whereby mV was plotted against temperature, was calculated for all individual TRIS calibrations separately. Second, the pH of TRIS was calculated with the sample’s temperature, using equation 5 of SOP 6a from Dickson et al., (2007). Third, pH_T_ of the sample was calculated using equation 8 of SOP 6a from Dickson et al., (2007). Finally, the respective calibration was applied to the mV raw data of the corresponding time for a given logger.

Dissolved oxygen – Before deployment, the loggers were calibrated using the lab calibration tool in the HOBO software. The loggers were placed in 100% saturated saltwater, whereafter they were placed in 0% saturated saltwater. 0% saturated saltwater was achieved by adding sodium sulphite. Dissolved oxygen data provided by the loggers were corrected in the HOBOware Dissolved Oxygen assistant with site-specific salinity data. If salinity data was not available for certain DO measurements, as the conductivity loggers had to be shuffled among sites and DO loggers did not, the average salinity of that site dataset was used. DO is provided in both mg/L and % saturation.

Temperature – Temperature was continuously measured by the conductivity, pH and DO loggers; however, only the temperature data from the pH loggers were used for the analysis of the sea surface temperature (SST) because the sample’s temperature was also used for the equation 5 of SOP 6a from Dickson et al., (2007). The loggers from the brand ‘HOBO’ give an accuracy of 0.2 °C for the temperature sensors whereas the loggers from the brand Odyssey do not.

Photosynthetically active radiation (PAR) – The Odyssey loggers have a planar cosine-corrected photosynthetic irradiance sensor (400 nm – 700 nm). This PAR sensor continually detects light intensity over a given time and records the summed value at the end of the logging period. To convert the sensor output to µmol m^-2^ s^-1^, all PAR loggers were deployed together with a Hydrolab DS5 Sonde (OTT Messtechnik GmbH & Co., Germany) equipped with a Li-Cor 4 pi spherical sensor, in Piscadera Bay Reef in front of the Carmabi research station (Carmabi Buoy 0: 12°07.417′ N, 68°58.592′ W) for 12 hours (7 am – 7 pm). Since it was not feasible to deploy the Hydrolab together with all PAR loggers at every study site, a new site was selected, Piscadera Bay Reef, which was in terms of benthos similar to the inland bays and the reference reefs. The PAR sensors of the loggers were deployed at the same height as the Li-Cor sensor of the Hydrolab with a logger interval of 15 minutes at a depth of 6 m. The deployment in Piscadera Bay Reef was needed for careful sensor calibration (Long et al. 2012). After retrieval of all PAR loggers and Hydrolab, the sensor output of the Odyssey loggers was calibrated with the Hydrolab data using a linear regression with the intercept forced through zero.

#### **Seawater carbonate chemistry, nutrients and trace metals**

High-resolution temporal monitoring of total alkalinity (TA) and nutrient concentrations was conducted during the dry season at each site for one day. In contrast, in the wet season, Spaanse Water Bay and Spaanse Water Reef were each monitored for one day, while Santa Martha Bay and Santa Martha Reef were monitored for two days, with water samples collected every 3-4 hours from sunrise until sunset. Additional discrete water samples were taken opportunistically throughout the fieldwork to cover the full range of weather conditions during the dry and wet season.

Total alkalinity and carbonate chemistry - Three 60 mL bottles were filled as close as possible to the monitoring stations and an equally divided mixture of seawater was made from these three bottles. Seawater was then filtered on site with a syringe filter of 0.2 µm (VWR) and a subsample was taken in a 60ml bottle. For each subsample the pH_T_, salinity and temperature were measured using a HACH multimeter. The subsamples were transported in a cooler to the research station, where they were poisoned with mercury chloride and stored in the dark at 4 °C until transportation to the University of Amsterdam (Amsterdam, the Netherlands) (ambient temperature during transport). At the University of Amsterdam, the water samples were stored again in the dark at 4 °C until further analysis.

Water samples for alkalinity were analysed within 3 months (cooler dry season) and within 1 month (warmer wet season). TA was analysed using potentiometric titration on a Metrohm 716 DMS Titrino (Herisau, Switzerland). TA was calculated according to standard protocols for seawater CO_2_ measurements (Dickson et al., 2007) using the at function in the seacarb package (version = 3.2.14) in R (2021.09.0) and the constants recommended by Dickson et al., (2007). pH, temperature and salinity values needed for this analysis were taken from measurements of the HACH multimeter which were taken immediately after water sample collection (see above). The accuracy and precision of the TA instruments were checked using certified reference material (CRM, batch 183 and 187) from the laboratory of A. Dickson (Scripps Institution of Oceanography). Accuracy was calculated as the average offset from certified values of TA. For the cooler dry season and the warmer wet season, TA values were within 17.26 μmol kg−1 (n = 13) and 54.70 μmol kg−1 (n = 24) of the certified value, respectively.

Seawater *p*CO_2_ and aragonite saturation state (Ω_Ar_) were calculated from in situ pH_T_, temperature, salinity and TA using the Carb function (flag=8) in the seacarb package in R.

Nutrient water samples - Three 60 mL bottles were filled as close as possible to the monitoring stations and an equally divided mixture of seawater was made from these three bottles. Seawater was then filtered on site with a syringe filter of 0.2 µm (VWR) and stored in 8 mL vials in a cooler during transport back to the research station, then at -20 °C until transportation to the University of Amsterdam on dry ice. At the University of Amsterdam, the water samples were stored again at -20 °C until further analysis. Nitrate, nitrite, ammonium and phosphate concentrations (µmol L^-1^) were measured using a Skalar SAN++ system autoanalyzer.

Seagrass samples - Seagrass samples were collected only from the inland bays because no seagrass was present at the reefs and were only collected during the wet season. Seagrass was collected from one sampling site that was as close as possible to the monitoring stations in the bays. Ten shoots (1 meter in between shoots) of *Thalassia testudinum* and *Halophila stipulacea* with roots were manually collected in Spaanse Water Bay, whereas only 10 shoots of *Halophila stipulacea* were collected in Santa Martha Bay due to the absence of *Thalassia testudinum*. The seagrass shoots were split into roots, rhizomes and leaves, after which only the leaves were used. Epiphytes were removed by rinsing material with seawater. Samples was excluded if the epiphytes could not be removed. Per bay, two leaves of every shoot of a seagrass species were pooled, resulting in a total of 3 samples of *T. testudinum* and *H. stipulacea* from Spaanse Water Bay and *H. stipulacea* from Santa Martha Bay. The samples were dried at 60°C for 32 hours. The dried seagrass was transported to the University of Amsterdam in zip lock bags, where it was analysed for trace metals concentrations (As, Cd, Co, Cr, Cu, Fe, Ni, P, Pb, Se, Zn) and ratios and percentage of carbon (C), nitrogen (N) and phosphate (P). Prior to the analysis, the dried seagrass was ground into a fine powder using mortar and pestle. To analyse the C and N content, 5-10 mg of solid seagrass powder from each sample (*in duplo*) was weighed in tin capsules using an analytical micro-balance and shaped into little round balls. A standard measurement of sulfanilic acid was used, which was also weighed into tin capsules (5-10 mg). These closed tin capsules were loaded into an automatic sampler, CN(S) elemental analyser. For the trace metal analysis, samples were digested in Teflon pressure tubes, using 100 mg of solid seagrass powder of each sample, 8 ml HNO_3_ and 2 ml HCL. After 1 hour of reaction time, the tubes were placed in a digestion microwave (program: 20 min 180 °C, 10 min 180 °C, 200 min 70 °C). The samples were diluted to 50 mL with Milli Q water. The total concentrations of trace metals (As, Cd, Co, Cr, Cu, Fe, Ni, P, Pb, Se, Zn) in seagrass leave tissue were measured by emission spectrometry.

#### **Sediment traps**

To assess sediment settlement, two sediment traps per site were deployed “in the wet season 2020” to collect sediment particles in the water column during 16 and 19 days at all four sites (Table S1). Sediment traps were only deployed in the wet season (November 2020). Each trap consisted of a PVC plastic container (diameter of 6.5 cm by 30.5 cm high) and had a mesh inside at 7.5 cm deep from the trap mouth in order to hold back fish. Two traps per site, separated 85-100 cm from each other, were deployed near the monitoring station attached to a metal rod and the bottom of the traps were positioned 10 cm above the seafloor, which resulted in a height of 40 cm from the trap mouth above the substrate (Storlazzi et al., 2011). To determine if trapped sediments were from local resuspension or transportation, benthic sediment at the base of each trap was also sampled. After retrieval of the sediment traps, the content was poured into aluminium containers after which it rested for 24 hours in order for the sediment to sink down. The excess water was removed with a syringe and the sediment was poured into a smaller aluminium container, which was dried in the oven at 60 °C for 48 hours. Finally, the sediment was stored in 50 mL Greiner tubes until further analysis at the University of Amsterdam.

Sediment samples were analysed for weight and particle size characteristics. The dried sediment was resuspended in milli-Q water to allow salt to be dissolved and removed from the sample, which was done by centrifuging the resuspended sample at 2500 rpm, removing the excess water and adding new milli-Q water until the salt content was undeterminable by the salt meter. The washed sediments were oven-dried at 60 °C for 1 week, after which the sediment samples were separated into six different fragments (>1 mm, 500 to <1000 μm, 250 to <500 μm, 125 to <250 μm, 63 to <125 μm and <63 μm) by placing the sieve stack on a mechanical shaker for 5 minutes. Each of the sediment fractions was weighed to the nearest 0.001 g. Sediment trap collection rate (mg cm^2^ day^−1^) was calculated as the weight of sediment trapped (mg) divided by the number of days the trap was deployed and the surface area of the trap (cm^2^). The amount of organic matter was determined by burning the sediment samples in the muffle furnace at 450°C for 6 hours, after which the samples were weighed again to the nearest 0.001 g. The difference of weight before and after burning is the amount of organic matter and the organic matter trap collection rate was calculated in the same way as the sedimentation collection rate.

#### **Effect-based chemical water quality assessment**

Passive sampling devices – The pharmaceutical passive polar organic chemical integrative samplers (POCIS) configuration, containing hydrophilic-lipophilic balance (HLB) sorbent was applied for the sampling of a wide range of organic contaminants from the water (Alvarez et al. 2004). The HLB sorbent was obtained from the manufacturer (Waters) in 6 g solid-phase extraction (SPE) cartridges. The sorbent was conditioned in the original SPE column by sequentially eluting with 150 mL acetone, 150 mL dichloromethane and 150 mL methanol (all chromatography grade) and was dried under vacuum. The dried sorbent was stored in a clean glass container until POCIS assembly. Stainless steel rings (an inner diameter of 5.4 cm), nuts, bolts, and tools (trays, tweezers and spatula) were cleaned with chromatography grade acetone before the assembly of the samplers. The POCIS were constructed at the University of Amsterdam according to De Baat et al., 2020. Per POCIS, 200 mg HLB sorbent (Oasis HLB) was enclosed between two polyethersulfone (PES) diffusion limiting membrane filters (Pall Corporation; 0.1 μm pore size, 90 mm diameter). The two filter membranes containing the sorbent were sandwiched between two metal rings and closed securely with three sets of nuts and bolts (Fig. S2). After assembly, the POCIS were stored at 4°C in food-grade Mylar zip lock bags until deployment.

Deployment of passive samplers – Forty POCIS were transported to Curaçao without refrigeration and stored at 4 °C upon arrival until the field deployment. At each site, seven POCIS were attached to a stainless steel frame and retained in a stainless steel cage. The cage was placed on a cement block and deployed at the monitoring stations next to the cement block with loggers (Fig. 1b). In addition, seven unexposed POCIS were included as blanks, which were stored the entire field trip in the fridge at 4 °C.

Retrieval of passive samplers – The cages with the samplers were retrieved after 29 and 30 days from Santa Martha and Spaanse Water, respectively (Table S1) and each POCIS was stored in a food-grade Mylar zip lock bag. In the laboratory, each POCIS was disassembled and the HLB sorbent of all 7 POCIS per site was collected in one 50 mL Greiner tube. The tubes were subsequently stored at -20 °C until transportation to the University of Amsterdam on dry ice and were stored again at -20 °C upon arrival until extraction.

Extraction of HLB sorbent – The frozen HLB sorbent in the Greiner tubes was freeze-dried for 3 days at -52 °C in a Scanvac CoolSafe freeze-dryer. All equipment used for the extraction procedure was cleaned with 1:1 (v:v) solution of chromatography grade acetone and HPLC grade methanol (AC:ME). The recovered HLB sorbent was transferred to glass solid-phase extraction (SPE) columns (Supelco) and placed on a SPE manifold, whereafter the sorbent was eluted two times with 5 mL acetonitrile and methanol (AC:ME) (50% v/v) solvent under vacuum. The POCIS extracts were topped up to exactly 10 mL with AC:ME solvent by weight on the analytical balance, and stored at -20 °C until further analyses.

Calculation of sampled water volumes by POCIS – A sampling rate for POCIS of 0.1 L d^-1^ per sampler was used to determine the sampled volume at each location (Harman et al., 2012). A sampled volume is necessary to determine the concentration factor of the deployed POCIS to ensure comparison of bioassay responses between sites, and the use of a single average sampled volume is warranted when using adsorption sampler like POCIS (de Weert et al. 2020) . The calculation of sampled water volumes by POCIS was done according to De Baat et al., (2020). Recovered HLB sorbent was corrected for loss during retrieval and extraction. The fraction of the recovered sorbent was calculated by dividing the recovered sorbent mass by the initial sorbent mass (1.4 g for 7 POCIS). Finally, the sampled volume was determined by multiplying the fraction of the recovered sorbent by the estimated sampling volume per location and by the number of deployment days per location (21 L for 7 POCIS for Santa Martha and 20.3 L for 7 POCIS for Spaanse Water).

The relative enrichment of the POCIS extracts – To determine the relative enrichment of the POCIS extracts (L/mL), the sampled volume by POCIS (L) was divided by the volume of the extracts (mL). This concentration is required to compare bioassay responses between sites and to effect-based trigger values.

Sample pre-treatment – Before application in the different bioassays, the solvent of the POCIS extracts was exchanged for dimethyl sulfoxide (DMSO). This was done by evaporating the extracts until dryness under N_2_ flow at room temperature and redissolving in DMSO (Table S2).

#### ***In vivo* bioassays**

Algal bioassay - *Dunaliella tertiolecta* were batch-cultured in 500 mL glass Erlenmeyers on 150 mL A* medium (Table S3) at 120 rpm at room temperature. Light was provided 24 hours per day (30 µmol photons m^-1^ s^-1^). Prior to the bioassay, the cell density of *Dunaliella tertiolecta* was measured by counting cells in a Bücker hemocytometer under a light microscope. Cells located at the top and right line of the smaller squares were excluded if the cells were positioned and/or touched those lines.

*Dunaliella tertiolecta* was exposed to dilution series of the POCIS extract in black PP 96-wells plates (Greiner Bio-One) with a final total volume of 280 µL per well and each well had a final cell density of 1 x 10^5^ cells mL^-1^. The POCIS extracts were step-wise diluted in the 96-well plate (Table S4) to determine the concentration causing 50% photosynthetic inhibition (EC_50_). The incubations occurred under continuous actinic LED light (659 nm, ~45 μmol m^−2^ s^−1^) at room temperature. The effective photosystem II efficiency (ΦPSII) of the algal suspension was measured after 4.5 hours with a WATER-PAM (Walz, Germany).

The POCIS extracts were step-wise diluted in the 96-well plate to get a concentration series (n=8) of extract (100%, 30%, 10%, 3%, 1%, 0.3% and 0.1%) to determine the concentration causing 50% photosynthetic inhibition (EC_50_). The final volume of each well was 280 µL of which 10% was always algal culture with a final cell density of 1e^5^ cells/mL in each well of the 96 wells plate, 90% or less was A* medium, a maximum of 1% was extract (for the pipetting scheme, see Table S4). A positive atrazine control and a negative control in A* medium were included as well. The DMSO concentration never exceeded 1%. The addition of 28 μL of algae initiated the exposure and the effective photosystem II efficiency (ΦPSII) of the algal suspension was measured after 4.5 h with a WATER-PAM (Walz, Germany).

Minimum and maximum fluorescence were determined and ΦPSII was calculated as F_max_-F_min_/F_max_, whereafter this was transformed to percentages of the control. The bioassay was considered valid when the positive control caused photosynthetic inhibition of approximately 50-65% in comparison to the negative control.

Bacterial luminescence inhibition assay - The *Alliivibrio fischeri* bioluminescence assay was performed at the Vrije Universiteit Amsterdam and according to Hamers et al., 2001. The concentration series (n=2) used for this test was 100%, 30%, 10%, 3%, 1%. The final volume of each well was 200 µL 0.5% of which was extract, 44.5% was 2% NaCl solution and 50% was marine bacteria culture. Triclosan was used as a positive control in this test. Luminescence was measured after 15 min and 30 min exposure to the extracts.

Data analysis of *in vivo* bioassays – Dose-response curves were made in GraphPad Prism (GraphPad Software Inc., version 9.1.2.226, San Diego, CA, USA) and the EC_50_ was determined by fitting a non-linear regression on the data with a built-in log-logistic model. The non-linear regression was forced to have a hillslope of 1 and the top plateau was forced to be 100% and the bottom plateau 0% of signal of the respective bioassay. Toxicity was expressed in toxic units (TU) and was calculated by 1/EC_50_. Finally, the bioassays responses were corrected for the estimated sampled water volume of the passive samplers.

### **Supplemental Results**

### **Seawater carbonate chemistry**

Total alkalinity (TA) ranged between 2064 and 2474 μmol kg^-1^. Significantly higher (5.1%) TA levels were measured during the dry season compared to the wet season (p = <0.001, Table S12, Figure S5a). The pCO_2_ reached concentrations between 281 and 534 μatm, with ~7.1% lower average values found in Spaanse Water sites during the wet season compared to Santa Martha sites; however, no significant difference was found between either habitats or seasons (Table S12). Aragonite saturation state (Ω_Ar_) varied between 3.0 and 4.4 (Figure S5d), with a significant difference between seasons (p ≤ 0.001) in which the dry season had 9.6% higher values than the wet season (p ≤ 0.001).

#### **Seagrasses as indicators of nutrient and trace metal pollution**

Seagrass samples – In contrast to the water samples, the seagrass samples were only used to examine bay-specific location effects as seagrass was only found in the bays. The seagrass leaves of *H. stipulacea* from Santa Martha Bay had 0.48% and 8.61% lower C and N content, respectively, and 5.56% higher P content than leaves of the same species from Spaanse Water Bay (Table S10). *T. testudinium*, which was only found in Spaanse Water Bay, had higher C and N content and lower P content than *H. stipulacea* found in both bays. In Spaanse Water Bay, *T. testudinium* had 12.11% and 29.67% higher C and N content, respectively, and 26.26% lower P content than the seagrass leaves of *H. stipulacea*. Overall, higher C and N content and lower P content was found in Spaanse Water Bay in comparison to Santa Martha Bay, according to two different seagrass species. *Halophila stipulacea* and *T. testudinium* tended to show similar nutrient trends when collected from the same site. This was also observed in the C:N:P ratios (Table S10). The lowest C:N ratio and highest N:P ratio and C:P ratios were found in *T. testudinium* from Spaanse Water Bay. *Halophila stipulacea* from Santa Martha Bay showed the highest C:N ratio and the lowest C:N ratio and N:P ratio.

Trace metals – Seagrass leaves collected from both Santa Martha Bay and Spaanse Water Bay contained copper (Cu), iron (Fe), nickel (Ni) and zinc (Zn), whereas arsenic (As), cadmium (Cd), cobalt (Co), lead (Pb) and selenium (Se) were not detected (Table S11). In addition to these detected metals, chromium (Cr) was found in *H. stipulacea* from Santa Martha Bay (Table S11). The highest leaf metal concentrations for Cu, Ni and Zn were found in *T. testidinum* (collected in Spaanse Water only), while *H. stipulacea* from Santa Martha Bay had the highest Fe concentration. Overall, the trace metal concentrations tended to be higher in Spaanse Water Bay than Santa Martha Bay, which was also found in the similar trace metal trend of *T. testidinum* and *H. stipulacea* collected from Spaanse Water Bay, with the exception of Ni.

| **Supplementary Tables**  Table S1. Overview of deployment and retrieval of loggers for environmental variables and pollution. Dates are indicated as DD-MM-YYYY. | | | | | | | | | | | | | | |
| --- | --- | --- | --- | --- | --- | --- | --- | --- | --- | --- | --- | --- | --- | --- |
| **Site** |  | | **Abiotic parameter** | | | | | | | | | | | |
|  |  |  | Conductivity & Temperature | | pH | | Dissolved oxygen | | Photosynthetically active radiation | | Organic chemical pollution | | Sediment traps | |
|  | Season | | Dry | Wet | Dry | Wet | Dry | Wet | Dry | Wet | Dry | Wet | Dry | Wet |
| **Santa Martha Bay** | Deployment | | 06-03-2020 | 4-11-2020 | 12-03-2020 | 29-10-2020 | 06-03-2020 | 29-10-2020 | 06-03-2020 | 4-11-2020 &  26-11-2020 |  | 29-10-2020 |  | 11-11-2020 |
|  | Final retrieval | | 21-3-2020 | 23-11-2020 | 21-03-2020 | 01-12-2020 | 21-03-2020 | 01-12-2020 | 18-03-2020 | 01-12-2020 |  | 27-11-2020 |  | 27-11-2020 |
|  | Number of days logged | | 16 | 20 | 10 | 34 | 16 | 34 | 13 | 17 |  | 30 |  | 16 |
| **Santa Martha Reef** | Deployment | | 08-03-2020 & | 23-11-2020 | 12-03-2020 & | 29-10-2020 | 08-03-2020 | 29-10-2020 | 08-03-2020 &  18-03-2020 | 04-11-2020 &  26-11-2020 |  | 29-10-2020 |  | 11-11-2020 |
|  | Final retrieval | | 21-03-2020 | 01-12-2020 | 21-03-2020 | 01-12-2020 | 21-03-2020 | 01-12-2020 | 18-03-2020 | 01-12-2020 |  | 27-11-2020 |  | 27-11-2020 |
|  | Number of days logged | | 14 | 9 | 10 | 34 | 18 | 34 | 9 | 17 |  | 30 |  | 16 |
| **Spaanse Water Bay** | Deployment | | 07-03-2020 | 28-10-2020 | 13-03-2020 | 28-10-2020 | 07-03-2020 | 28-10-2020 | 07-03-2020 | 28-10-2020 &  17-11-2020 &  24-11-2020 |  | 30-10-2020 |  | 9-11-2020 |
|  | Final retrieval | | 21-03-2020 | 21-11-2020 | 21-03-2020 | 02-12-2020 | 21-03-2020 | 02-12-2020 | 19-03-2020 | 02-12-2020 |  | 27-11-2020 |  | 28-11-2020 |
|  | Number of days logged | | 15 | 25 | 9 | 36 | 15 | 36 | 13 | 23 |  | 29 |  | 19 |
| **Spaanse Water Reef** | Deployment | | 10-03-2020 | 21-11-2020 | 14-03-2020 | 28-10-2020 | 14-03-2020 | 28-10-2020 | 10-03-2020 | 30-10-2020 |  | 30-10-2020 |  | 9-11-2020 |
|  | Final retrieval | | 21-03-2020 | 02-12-2020 | 20-03-2020 | 02-12-2020 | 21-03-2020 | 02-12-2020 | 19-03-2020 | 14-11-2020 |  | 27-11-2020 |  | 28-11-2020 |
|  | Number of days logged | | 12 | 12 | 8 | 36 | 8 | 36 | 10 | 16 |  | 29 |  | 19 |

| Table S2. Overview of volumes used for evaporation of POCIS extracts dissolved in AC:ME solvent and redissolvement of the extracts in DMSO. | | | | |
| --- | --- | --- | --- | --- |
|  | **Bioassay** | **Extract volume for evaporation** | **DMSO volume for dissolving** | **Final enrichment** |
| *in vivo* | Bacterial bioluminescence inhibition | 1 mL | 25 µL | 40 x |
|  | Algal | 3 mL | 75 µL | 40 x |
| *in vitro* | PAH | 2 mL | 100 µL | 20 x |
|  | ERα |  |  |  |

| Table S3. Recipe for A* medium. *Dunaliella tertiolecta* were batch-cultured in glass Erlenmeyers on A* medium. directions for the recipe are as following: (1) Pre-rinse bottle with 10% HCL then several time with DI water; (2) Fill bottle ¾ with milli Q water (750 mL for 1L); (3) Weigh out **1 –** **2** and add. Mix until salts dissolved; (4) Add **3** **–** **7**; (5) Autoclave for 20 min at 121 °C; (6) Cool to room temperature and add **8** (stored in walk-in freezer), **9** and **10** (stored in walk-in freezer) under hood. Keep **8 – 10** sterile. | | | | |
| --- | --- | --- | --- | --- |
| **Step** | Compound | | Addition to 1L | |
| **1** | MgSO_4_•7H_2_O – M33 | | 5 g | |
| **2** | NaCl | | 25 g | |
| **3** | Salt stock | | 25 mL | |
| **4** | Na_2_SiO_3_•5H_2_O stock | | 10 mL | |
| **5** | H_3_BO_3_ stock | | 10 mL | |
| **6** | Metals working stock | | 10 mL | |
| **7** | NaNO_3_ | | 10 mL | |
| After autoclaving | | | | |
| **8** | Vitamins working stock | | 1 mL | |
| **9** | K_2_HPO_4_•3H_2_O | | 1 mL | |
| **10** | NaHCO_3_ | | 1 mL | |
| Table S4. Pipetting scheme for PAM-test with marine algae *(Dunaliella tertiolecta)*. One 96-wells plate, consisting of 12 columns and 8 rows, corresponded to one specific extract as a concentration series was made. The dilutions were made by pipetting 28 uL of the highest concentration (100% and 30%) into a subsequential lower concentration and so on until the lowest concentration was reached (0.1% and 0.3%), whereafter the last 28 uL was transferred to the waste wells in column 12. This pipetting scheme below was retrieved from Bas van Beusekom (2018) and corrected for this specific test with the marine algae. | | | | |
| A* medium | | Column 1: Blanco | | 280 μL |
|  |  | Column 2: Fluorescence control | | 279 μL |
|  |  | Column 9: Low concentration | | 279 μL |
|  |  | Column 10: High concentration | | 277 μL |
|  |  | Column 3: Negative control | | 252 μL |
|  |  | Column 4 - 8: Experiment | | 252 μL |
|  |  | Column 11: Atrazine control | | 249 μL |
| DMSO | | Column 2: Fluorescence control | | 0.9 μL |
| POCIS extract | | Column 10: High concentration | | 2.8 μL |
|  |  | Column 9: Low concentration | | 0.9 μL |
| Dilutions and transfers | | High: Column 10 (100%) 🡪 Column 8 (10%) 🡪 Column 6 ( 1%) 🡪 Column 4 (0.1%) 🡪 Column 12 (waste) | | 28 μL |
|  | | Low: Column 9 (30%) 🡪 Column 7 (3%) 🡪 Column 5 (0.3%) 🡪 Column 12 (waste) | | 28 μL |
| Atrazine | | Column 11: Atrazine control | | 2.8 μL |
| Marine algae | | Column: 3 t/m 11 | | 28 μL |

| Table S5. Scleractinian coral diversity described for all four sites. | | | | |
| --- | --- | --- | --- | --- |
| **Coral species** | **Sites** | | | |
|  | Santa Martha Bay | Santa Martha Reef | Spaanse Water Bay | Spaanse Water Reef |
| *Agaricia agaricites* |  | X |  |  |
| *Diploria clivosa* |  |  |  | X |
| *Diploria labyrinthiformis* |  | X |  |  |
| *Diploria strigosa* |  | X |  | X |
| *Favia fragum* | X |  |  |  |
| *Madracis mirabilis* |  |  |  | X |
| *Millepora alcicornis* |  |  |  | X |
| *Millepora complanata* |  | X |  | X |
| *Millepora* sp. |  |  |  | X |
| *Montastraea annularis* |  | X |  | X |
| *Montastraea cavernosa* |  |  |  | X |
| *Porites astreoides* |  | X |  | X |
| *Porites porites* | X |  |  |  |
| *Siderastrea radians* | X |  | X |  |
| *Siderastrea siderea* | X | X | X | X |
| *Stephanocoenia michelini* |  | X |  |  |
| Unknown |  |  |  | X |
|  | Total n = 4 | Total n = 9 | Total n = 2 | Total n = 12 |

| Table S6*.* Physicochemical parameters for all four sites studied in March 2020 (cool, dry season) and Oct/Nov 2020 (warm, wet season). SE = standard error and CV = coefficient of variation. DO = dissolved oxygen, PAR = photosynthetically active radiation. | | | | | | | | | | | | |
| --- | --- | --- | --- | --- | --- | --- | --- | --- | --- | --- | --- | --- |
| **Site** |  | | **Environmental parameter** | | | | | | | | | |
|  |  |  | Temperature (°C) | | Salinity (ppt) | | pH (total scale) | | DO (mg L^-1^) | | PAR (µmol m^-2^ s^-1^) | |
|  | Season | | Dry | Wet | Dry | Wet | Dry | Wet | Dry | Wet | Dry | Wet |
| **Santa Martha Bay** | Mean (SE) | | 27.5 (0.013) | 28.9 (0.013) | 35.6 (0.009) | 33.7 (0.038) | 8.05 (0.001) | 8.08 (0.001) | 6.09 (0.006) | 6.30 (0.016) | 559 (10.4) | 360 (7.9) |
|  | Range (CV) | | 2.7 (0.024) | 3.5 (0.020) | 3.2 (0.016) | 7.8 (0.048) | 0.09 (0.002) | 0.34 (0.006) | 2.42 (0.059) | 6.52 (0.144) |  |  |
|  | Daily max. range | | 2.57 | 2.0 | 2.9 | 6.5 | 0.05 | 0.22 | 2.34 | 5.41 |  |  |
|  | Max. | | 29.3 | 30.4 | 37.3 | 35.7 | 8.08 | 8.23 | 7.16 | 9.1 | 1644 | 1544 |
|  | Min. | | 26.6 | 26.9 | 34.1 | 27.8 | 7.99 | 7.89 | 4.74 | 2.58 |  |  |
| **Santa Martha Reef** | Mean (SE) | | 27.0 (0.003) | 28.9 (0.004) | 35.7 (0.005) | 33.7 (0.012) | 8.03 (0.001) | 8.04 (0.001) | 6.23 (0.005) | 6.19 (0.007) | 813 (17.1) | 412 (12.9) |
|  | Range (CV) | | 1.0 (0.006) | 1.1 (0.010) | 1.7 (0.008) | 1.4 (0.010) | 0.07 (0.002) | 0.25 (0.006) | 1.5 (0.053) | 2.84 (0.063) |  |  |
|  | Daily max. range | | 0.8 | 0.7 | 1.0 | 0.9 | 0.06 | 0.10 | 1.43 | 2.0 |  |  |
|  | Max. | | 27.5 | 29.5 | 36.5 | 34.2 | 8.07 | 8.20 | 6.95 | 7.58 | 2122 | 1153 |
|  | Min. | | 26.5 | 28.3 | 34.9 | 32.9 | 8.00 | 7.95 | 5.46 | 4.74 |  |  |
| **Spaanse Water Bay** | Mean (SE) | | 27.8 (0.008) | 29.3 (0.007) | 35.5 (0.019) | 35.3 (0.018) | 8.05 (0.002) | 7.95 (0.001) | 6.38 (0.012) | 5.75 (0.016) | 351 (7.3) | 275 (6.5) |
|  | Range (CV) | | 2.0 (0.014) | 2.5 (0.017) | 3.3 (0.032) | 3.1 (0.025) | 0.33 (0.011) | 0.33 (0.006) | 4.89 (0.114) | 6.45 (0.157) |  |  |
|  | Daily max. range | | 1.6 | 1.3 | 1.7 | 1.5 | 0.13 | 0.25 | 3.15 | 5.91 |  |  |
|  | Max. | | 28.9 | 30.4 | 36.9 | 36.6 | 8.19 | 8.11 | 8.61 | 9.38 | 1392 | 930 |
|  | Min. | | 26.9 | 27.9 | 33.6 | 33.5 | 7.87 | 7.78 | 3.72 | 2.93 |  |  |
| **Spaanse Water Reef** | Mean (SE) | | 27.0 (0.003) | 29.0 (0.005) | 35.5 (0.003) | 35.8 (0.007) | 8.04 (0.001) | 8.05 (0.001) | 6.52 (0.006) | 6.18 (0.009) | 591 (14.1) | 452 (14.4) |
|  | Range (CV) | | 0.7 (0.006) | 2.0 (0.012) | 1.2 (0.005) | 1.4 (0.006) | 0.10 (0.002) | 0.28 (0.006) | 1.86 (0.044) | 3.44 (0.081) |  |  |
|  | Daily max. range | | 0.6 | 0.8 | 1.1 | 1.0 | 0.09 | 0.20 | 1.66 | 3.14 |  |  |
|  | Max. | | 27.3 | 30.0 | 36.1 | 36.3 | 8.08 | 8.17 | 7.12 | 7.86 | 1759 | 1377 |
|  | Min. | | 26.6 | 28.0 | 34.9 | 34.9 | 7.98 | 7.89 | 5.26 | 4.42 |  |  |

| Table S7. Results from Scheirer-Ray-Hare extension of the Kruskal-Wallis test comparing daily averages of temperature, salinity, pH, dissolved oxygen (DO) and photosynthetically active radiation (PAR) across habitats and seasons. Df = Degrees of freedom, Sum sq =sum of squares and * indicates significant p-values (<0.05). | | | | | | | |
| --- | --- | --- | --- | --- | --- | --- | --- |
| **Environmental parameter**  (daily averages) |  | **Scheirer-Ray-Hare extension of the Kruskal-Wallis test** | | | | **Posthoc Dunn-test** | |
|  | Term | df | Sum Sq | H-value | p-value |  | p-value |
| Temperature (°C) | Habitat | 1 | 8657 | 4.083 | 0.043* | Bay > Reef | 0.0736 |
|  | Season | 1 | 202735 | 95.63 | <0.001* | Dry < Wet | <0.001* |
|  | Habitat:Season | 1 | 1239 | 0.584 | 0.445 |  |  |
| Salinity (ppt) | Habitat | 1 | 216 | 0.170 | 0.680 |  |  |
|  | Season | 1 | 15538 | 12.225 | 0.005* | Dry > Wet | <0.001* |
|  | Habitat:Season | 1 | 610 | 0.480 | 0.489 |  |  |
| pH (total scale) | Habitat | 1 | 7049 | 4.540 | 0.033* | Bay < Reef | 0.035* |
|  | Season | 1 | 809 | 0.521 | 0.470 |  |  |
|  | Habitat:Season | 1 | 3499 | 2.253 | 0.133 |  |  |
| DO (mg L^-1^) | Habitat | 1 | 28467 | 9.315 | 0.002* | Bay < Reef | 0.005* |
|  | Season | 1 | 42905 | 14.040 | <0.001* | Dry > Wet | <0.001* |
|  | Habitat:Season | 1 | 0 | 0.0001 | 0.993 |  |  |
| PAR (µmol m^-2^ s^-1^) | Habitat | 1 | 25129 | 17.982 | <0.001* | Bay < Reef | <0.001* |
|  | Season | 1 | 24247 | 17.422 | <0.001* | Dry > Wet | <0.001* |
|  | Habitat:Season | 1 | 4 | 0.003 | 0.957 |  |  |

| Table S8. Results from Scheirer-Ray-Hare extension of the Kruskal-Wallis test comparing daily variability of temperature, salinity, pH, dissolved oxygen (DO) and photosynthetically active radiation (PAR) across habitats and seasons. Df = Degrees of freedom, Sum sq =sum of squares and * indicates significant p-values (<0.05). | | | | | | | |
| --- | --- | --- | --- | --- | --- | --- | --- |
| **Environmental parameter**  (daily variability) |  | **Scheirer-Ray-Hare extension of the Kruskal-Wallis test** | | | | **Posthoc Dunn-test** | |
|  | Term | df | Sum Sq | H-value | p-value |  | p-value |
| Temperature (°C) | Habitat | 1 | 132536 | 74.123 | <0.001* | Bay > Reef | <0.001* |
|  | Season | 1 | 14718 | 8.231 | 0.004* | Dry < Wet | 0.003* |
|  | Habitat:Season | 1 | 74 | 0.041 | 0.839 |  |  |
| Salinty (ppt) | Habitat | 1 | 20723 | 16.304 | <0.001* | Bay > Reef | <0.001* |
|  | Season | 1 | 12350 | 9.717 | 0.002* | Dry > Wet | 0.010* |
|  | Habitat:Season | 1 | 6 | 0.005 | 0.944 |  |  |
| pH_T_ | Habitat | 1 | 21837 | 14.064 | <0.001* | Bay > Reef | <0.001* |
|  | Season | 1 | 32824 | 21.140 | <0.001* | Dry > Wet | <0.001* |
|  | Habitat:Season | 1 | 7290 | 4.6955 | 0.0302* | See text |  |
| DO (mg L^-1^) | Habitat | 1 | 200388 | 65.576 | <0.001* | Bay > Reef | <0.001* |
|  | Season | 1 | 107825 | 35.285 | <0.001* | Dry < Wet | <0.001* |
|  | Habitat:Season | 1 | 14706 | 4.183 | 0.028* | See text |  |
| PAR  (µmol m^-2^ s^-1^) | Habitat | 1 | 24277 | 17.3714 | <0.001* | Bay < Reef | <0.001* |
|  | Season | 1 | 42463 | 30.385 | <0.001* | Dry > Wet | <0.001* |
|  | Habitat:Season | 1 | 270 | 0.193 | 0.661 |  |  |

| Table S9. Results from Scheirer-Ray-Hare extension of the Kruskal-Wallis test comparing average nitrate, ammonium and phosphate concentrations of the water samples across habitat and season. Df = Degrees of freedom, Sum sq =sum of squares and * indicates significant p-values (<0.05). | | | | | | | |
| --- | --- | --- | --- | --- | --- | --- | --- |
| **Nutrients** |  | **Scheirer-Ray-Hare extension of the Kruskal-Wallis test** | | | | Posthoc Dunn-test | |
|  | **Term** | **df** | **Sum Sq** | **H-value** | **p-value** |  | p-value |
| Nitrate (NO_3_) | Habitat | 1 | 871 | 1.306 | 0.253 |  |  |
|  | Season | 1 | 31871 | 47.801 | 0* | Dry < Wet | <0.001* |
|  | Habitat:Season | 1 | 1057 | 1.585 | 0.208 |  |  |
| Ammonium (NH_4_) | Habitat | 1 | 2652 | 3.974 | 0.046* | Bay > Reef | 0.066 |
|  | Season | 1 | 34211 | 51.263 | 0* | Dry < Wet | <0.001* |
|  | Habitat:Season | 1 | 457 | 0.684 | 0.408 |  |  |
| Phosphate (PO_4_) | Habitat | 1 | 898 | 1.376 | 0.241 |  |  |
|  | Season | 1 | 9676 | 14.830 | <0.001* | Dry < Wet | <0.001* |
|  | Habitat:Season | 1 | 1054 | 1.616 | 0.204 |  |  |

| Table S10. Leaf nutrient concentrations of two seagrass species in the two inland bays collected in Oct/Nov 2020 (warm, wet season). Ratios are mol ratios. | | | | | | | | |
| --- | --- | --- | --- | --- | --- | --- | --- | --- |
| Bay | Seagrass species | % C | % N | % P | C:N | N:P | C:P | C:N:P |
| Santa Martha | *Halophila stipulacea* | 31.24 | 1.91 | 0.19 | 19 | 23 | 435 | 435:19:01 |
| Spaanse Water | *Halophila stipulacea* | 31.39 | 2.09 | 0.18 | 17 | 25 | 438 | 438:17:01 |
| Spaanse Water | *Thalassia testidinum* | 35.19 | 2.71 | 0.15 | 15 | 39 | 593 | 593:15:01 |

| Table S11. Trace metal concentrations in seagrass leaves (in µg g^-1^) of two seagrass species in two inland bays collected in Oct/Nov 2020 (warm, wet season). nd = not detected. | | | | | | | | | | | |
| --- | --- | --- | --- | --- | --- | --- | --- | --- | --- | --- | --- |
| Bay | Seagrass species | As | Cd | Co | Cr | Cu | Fe | Ni | Pb | Se | Zn |
| Santa Martha | *Halophila stipulacea* | nd | nd | nd | 16.02 | 13.78 | 1538.55 | 4.93 | nd | nd | 1.30 |
| Spaanse Water | *Halophila stipulacea* | nd | nd | nd | nd | 23.33 | 219.37 | 2.75 | nd | nd | 24.60 |
| Spaanse Water | *Thalassia testidinum* | nd | nd | nd | nd | 37.19 | 244.03 | 14.93 | nd | nd | 68.99 |

| Table S12. Results from Scheirer-Ray-Hare extension of the Kruskal-Wallis test comparing average *p*CO_2_, saturation state for aragonite (Ω_Ar_) and total alkalinity (TA) of the water samples across habitat and season. Df = Degrees of freedom, Sum sq =sum of squares and * indicates significant p-values (<0.05). | | | | | | | |
| --- | --- | --- | --- | --- | --- | --- | --- |
| **Nutrients** |  | **Scheirer-Ray-Hare extension of the Kruskal-Wallis test** | | | | Posthoc Dunn-test | |
|  | **Term** | **df** | **Sum Sq** | **H-value** | **p-value** |  | p-value |
| pCO_2_ | Habitat | 1 | 353 | 0.783 | 0.376 |  |  |
|  | Season | 1 | 166 | 0.369 | 0.543 |  |  |
|  | Habitat:Season | 1 | 26 | 0.057 | 0.811 |  |  |
| Ω_Ar_ | Habitat | 1 | 568.2 | 1.262 | 0.261 |  |  |
|  | Season | 1 | 7123.3 | 15.824 | <0.001* | Dry > Wet | <0.001* |
|  | Habitat:Season | 1 | 48.2 | 0.107 | 0.744 |  |  |
| TA | Habitat | 1 | 300 | 0.666 | 0.414 |  |  |
|  | Season | 1 | 20398.1 | 45.312 | <0.001* | Dry > Wet | <0.001* |
|  | Habitat:Season | 1 | 92.4 | 0.205 | 0.651 |  |  |

### **Supplementary Figures**

| 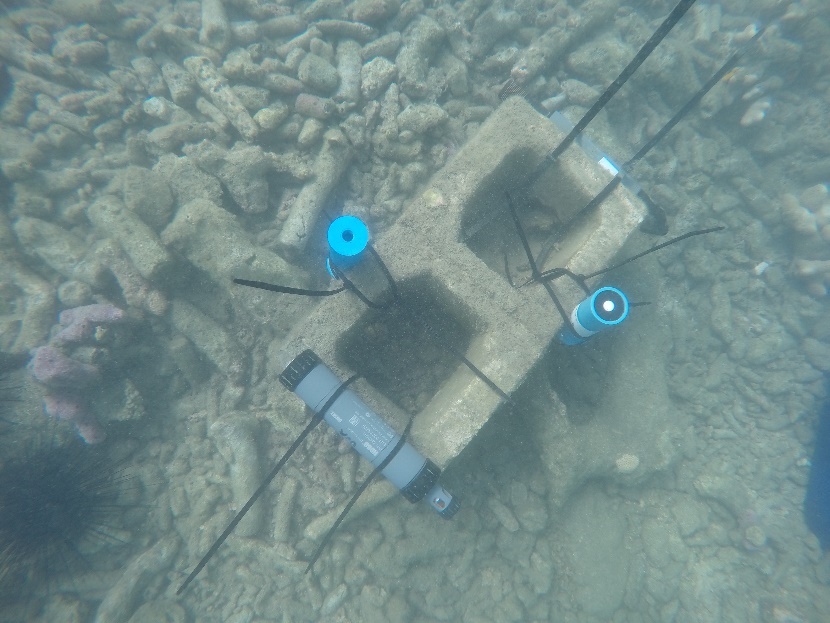  a | 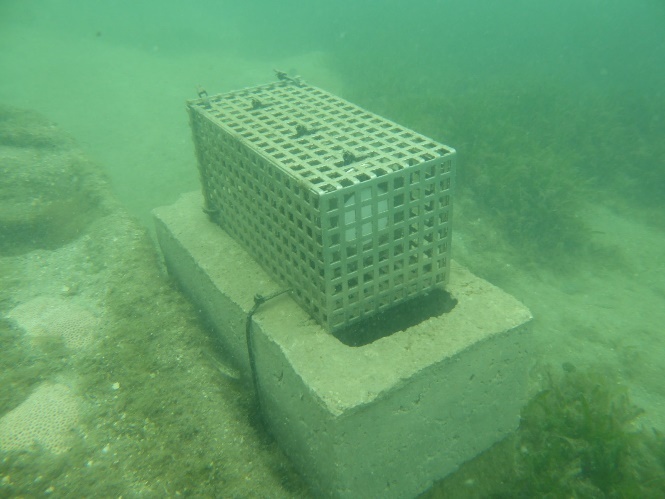  b |
| --- | --- |
| *Figure S1.* Cement block at the monitoring stations with (a) loggers and (b) passive samplers. Pictured at monitoring stations in the inland bays. | |

| 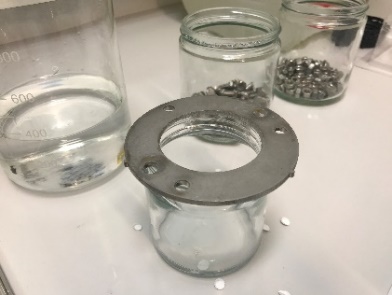 | 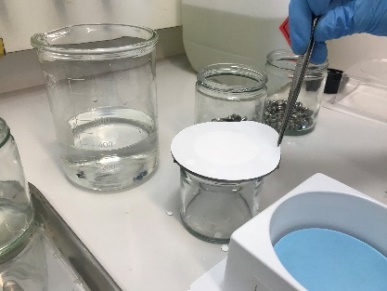 | 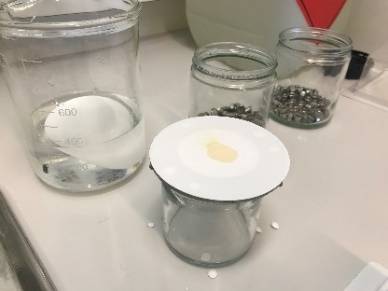 |
| --- | --- | --- |
| 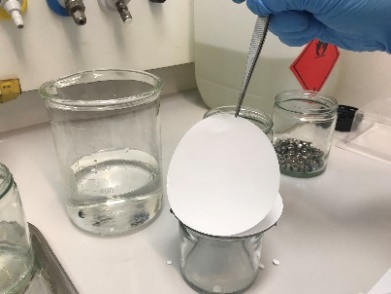 | 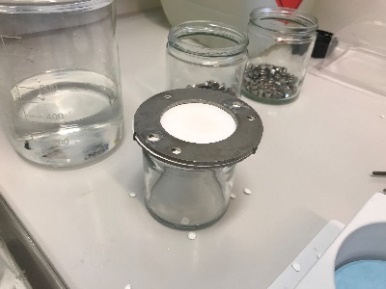 | 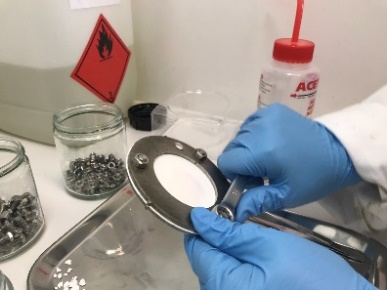 |
| *Figure S2.* Constructing POCIS passive sampling devices | | |

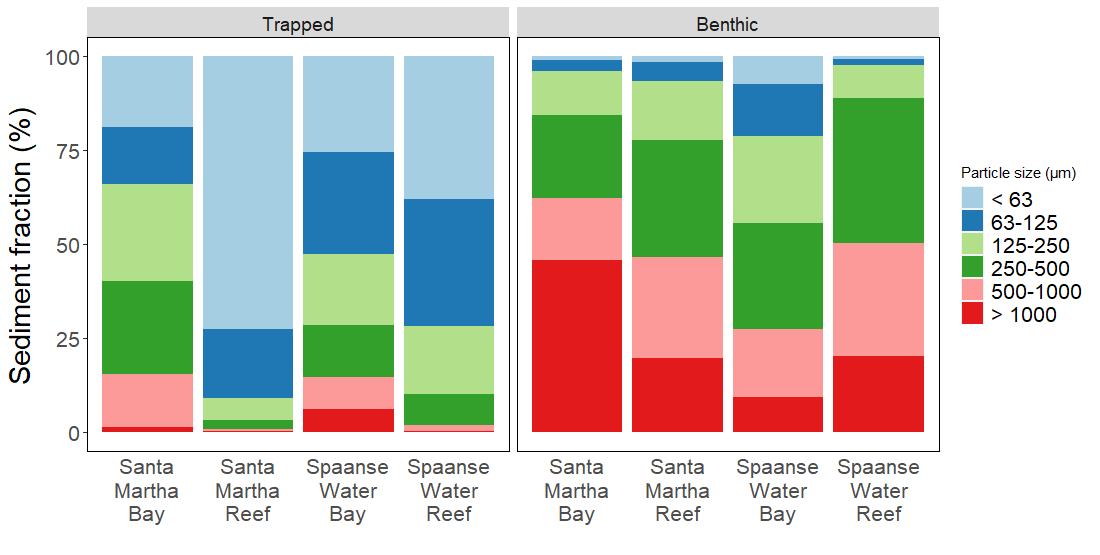

*Figure S3*. Particle size data from the sediment traps and the benthos at all four sites during the warm, wet season (Oct-Nov 2020) are shown in sediment fraction (%).

| 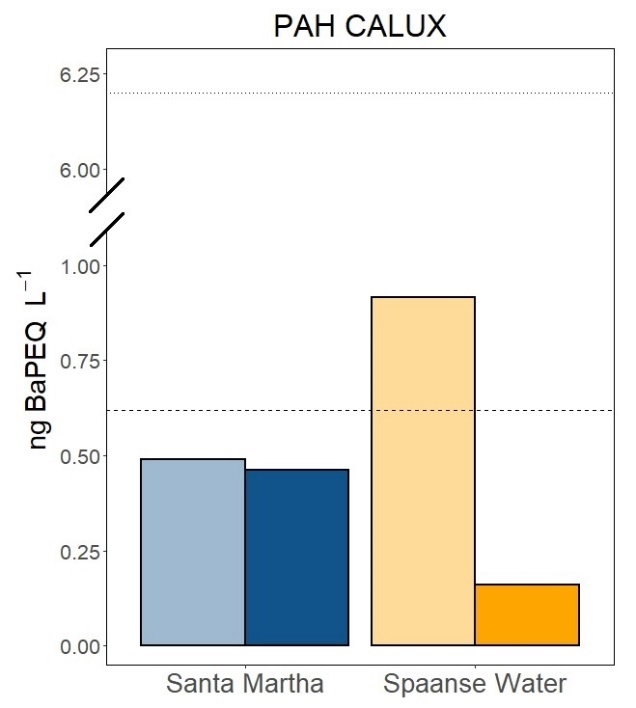a | 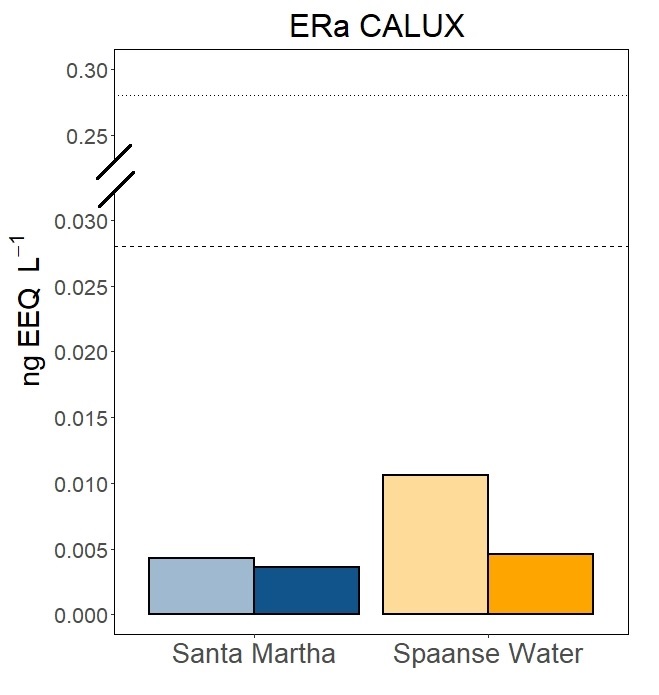b |
| --- | --- |
| 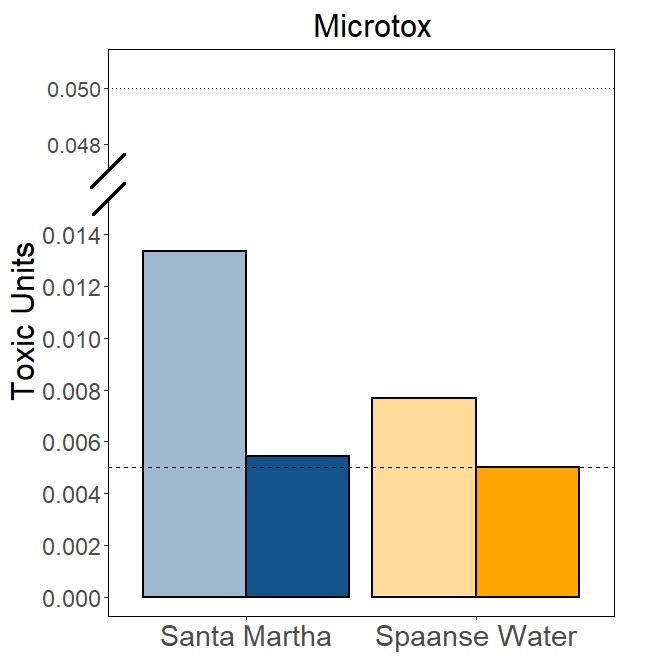c | 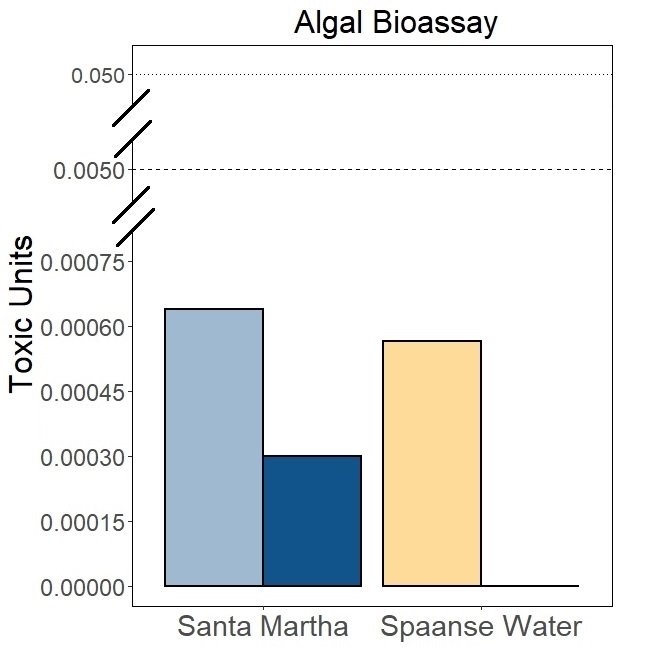d |
| 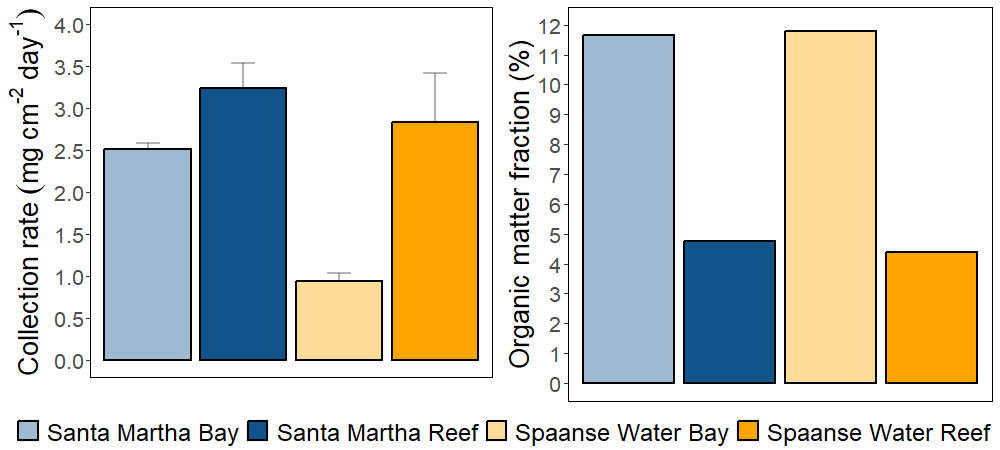 | |
| *Figure S4*. Reponses of (a) PAH CALUX, (b) ERα CALUX, (c) bacterial bioluminescence inhibition, and (d) Algal bioassays to extracts of POCIS from four surface water locations. Dotted line = effect-based trigger value (EBT) for potential ecotoxicological risks (freshwater) and dashed line = preliminary effect-based trigger (iEBT) value for potential ecotoxicological (marine) environment. Note different y-scales between graphs and scale breaks on y-axis. | |

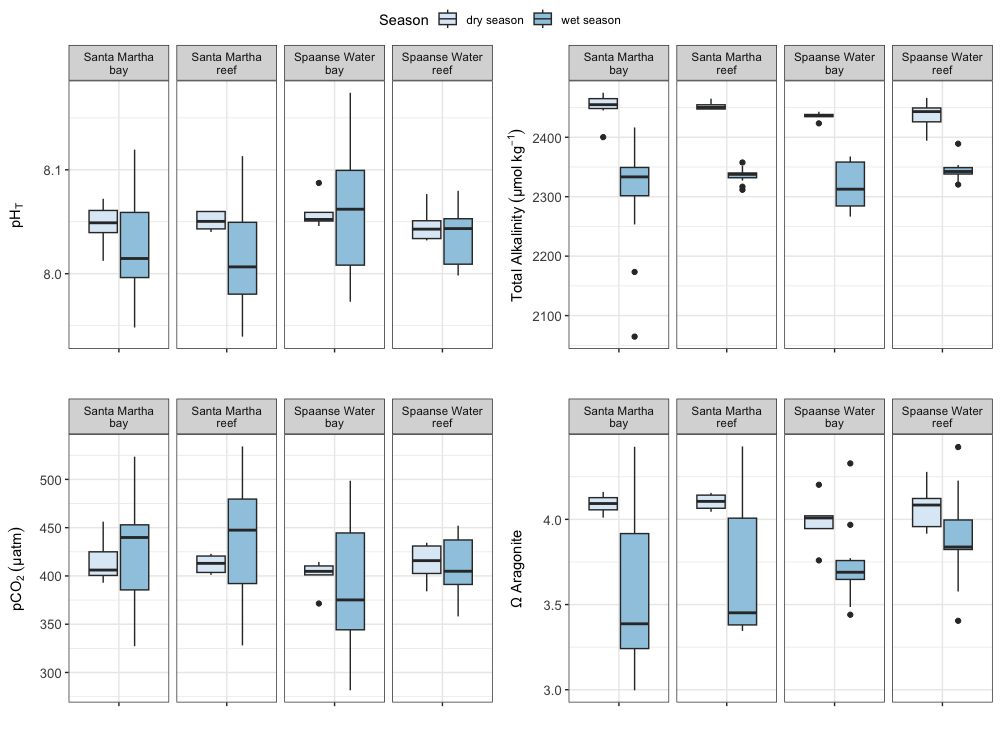

d

c

b

a

*Figure S5*. Boxplots for carbonate chemistry parameters of discrete water samples collected in March 2020 (cool, dry season) and Oct/Nov 2020 (warm, wet season). a. pH_T_, b. Total Alkalinity (TA) (μmol kg^-1^), c. pCO_2_ (μatm), d. Saturation state for aragonite (Ω_Ar_). Boxes span the ﬁrst to third quartiles; the horizontal line inside the boxes represents the median, black dots represent the outliers between the range of 3 times the inter-quartile range (IQR) as extreme outliers were excluded that exceeded 3 times the IQR. Whiskers represent the range of the data that falls within 1.5 times the IQR.

1. The conductivity of all solutions varies with temperature, however, the Odyssey Salinity program corrects for the temperature coefficient such that the conductivity output by the logger is the conductivity of the solution at 25 °C. The temperature output provided by the conductivity logger is only displayed for reference and does not have to be taken into account.
